# Rubisco is evolving for improved catalytic efficiency and CO_2_ assimilation in plants

**DOI:** 10.1101/2022.07.06.498985

**Authors:** Jacques W. Bouvier, David M. Emms, Steven Kelly

## Abstract

Rubisco is the primary entry point for carbon into the biosphere. However, rubisco is widely regarded as inefficient leading many to question whether the enzyme can adapt to become a better catalyst. Through a phylogenetic investigation of the molecular and kinetic evolution of Form I rubisco we demonstrate that rubisco is not stagnant. Instead, we demonstrate *rbcL* is among the 1% of slowest evolving genes and enzymes on Earth, accumulating one nucleotide substitution every 0.9 million years and one amino acid mutation every 7.2 million years. Despite this, we demonstrate that rubisco catalysis is continuing to evolve toward improved CO_2_/O_2_ specificity, carboxylase turnover, and carboxylation efficiency. Consistent with this kinetic adaptation, we reveal that increased rubisco evolution leads to a concomitant improvement in leaf-level CO_2_ assimilation. Thus, rubisco is continually evolving toward improved catalytic efficiency and CO_2_ assimilation in plants.

## Introduction

Ribulose-1,5-bisphosphate carboxylase/oxygenase (rubisco) converts atmospheric CO_2_ into sugars that fuel the majority of life on Earth. The enzyme evolved ∼3 billion years ago when the atmosphere contained high levels of CO_2_ (≥10,000% present atmospheric levels) and comparatively little O_2_ (≤0.1% present atmospheric levels) (Figure 1)^1–7^. Since emergence, the enzyme has helped guide the atmosphere on a trajectory of increasing O_2_ and declining CO_2_ (Figure 1)^1,8^ such that current concentrations of CO_2_ (0.04%) and O_2_ (20.95%) are inverted compared to when the enzyme first evolved (Figure 1).

**Figure 1.**
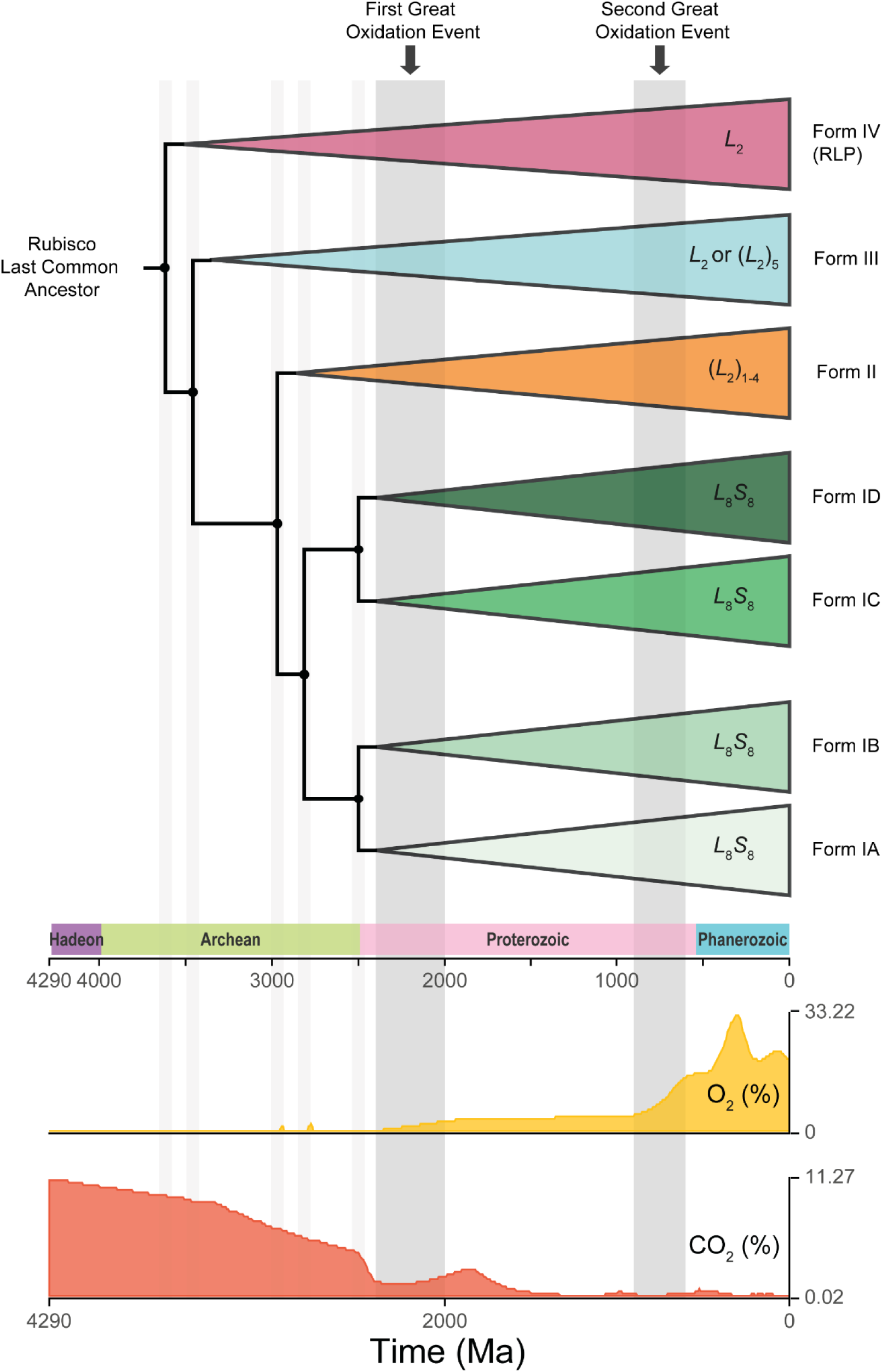
The evolutionary history of rubisco in the context of atmospheric CO_2_ (%) and O_2_ (%) following divergence from the ancestral rubisco-like protein (RLP). For ease of visualisation, branch points in the phylogeny are indicated by grey vertical bars. The First and Second Great Oxidation events are also indicated by grey vertical bars and have been labelled. Graphics of atmospheric CO_2_ and O_2_ levels were adapted from the *TimeTree* resource (http://www.timetree.org; ^84^).

Although all extant rubisco are descended from a single ancestral rubisco-like protein^9–11^, the enzyme is found in a variety of compositional forms across the tree of life (Figure 1)^12,13^. The simplest manifestations are the Form II and Form III variants found in protists, archaea, and some bacteria which are composed of a dimer, or dimers, of the ∼50 kDa rubisco large (RbcL) subunit^13–16^. In contrast, Form I rubisco is a hexadecamer comprised of four RbcL dimers organised in an antiparallel core capped at either end by the ∼15 kDa rubisco small subunit (RbcS)^13,17^. Of these three Forms, only Form I and II have been recruited for oxygenic photosynthesis^16^, with Form I being responsible for the vast majority of global CO_2_ assimilation^16,18^.

Within Form I rubisco the active site is located in RbcL^15,19,20^. As a result, interspecific differences in Form I kinetics are primarily attributable to sequence variation in RbcL^21–32^. Despite not playing a direct role in catalysis RbcS influences the function of rubisco^33^, and its incorporation in the holoenzyme enables its higher kinetic efficiency^34^. Specifically, RbcS enhances the stability and assembly of the holoenzyme complex^20,35–40^, improves the efficiency CO_2_ binding^41^, and is thought to act as a reservoir for CO_2_ accumulation^42^. Accordingly, rubisco function is altered when RbcS is mutated^43–45^, or when chimeric holoenzymes are created *in vivo*^46–50^ and *in vitro*^51–56^. Moreover, there is increasing recognition of the importance of both environment^57^ and organ-specific^58,59^ differences in plant RbcS isoform expression on holoenzyme catalysis. However, even though RbcS influences holoenzyme function, sequence variation in RbcL remains the primary determinant of variation in kinetics^21–32^.

Although there is kinetic variability between rubisco orthologs, the enzyme is considered to be an inefficient catalyst. For example, the maximum substrate-saturated turnover rate of Form I rubisco (<12 s^-1^)^60^ is slower than average^61^. In addition, rubisco catalyses a reaction with O_2_ ^62–64^ that is competitive with CO_2_ and results in the loss of fixed carbon via photorespiration^65–67^. As a consequence, rubisco appears poorly suited to the current O_2_-rich/CO_2_-poor atmosphere (Figure 1). Moreover, it appears that instead of improving enzyme function, multiple lineages have evolved alternative strategies to overcome rubisco’s shortcomings. For example, higher rates of CO_2_ assimilation are often achieved either by synthesising large quantities of rubisco (∼50% of soluble protein in plants^68^ and some microbes^69,70^), or by operating CO_2_-concentrating mechanisms^71–73^. As a result, many have questioned whether the enzyme is already perfectly adapted, and whether further kinetic improvements are possible^15,63,67,74–78^. Obtaining answers to these questions would shed light on the “rubisco paradox” – helping to explain why this enzyme of such paramount importance appears poorly adapted for its role.

The initial hypothesis that attempted to explain the above rubisco paradox proposed that rubisco is constrained by catalytic trade-offs that limit the enzyme’s adaptation. This theory was pioneered by two studies^79,80^ which found antagonistic correlations between rubisco kinetic traits and proposed that these trade-offs were caused by constraints on its catalytic mechanism. However, recent evidence has questioned this hypothesis as the sole mechanism to explain the rubisco paradox. Specifically, analysis of larger species sets have revealed that kinetic trait correlations are not strong^81–83^. In addition, phylogenetic signal in rubisco kinetics causes kinetic trait correlations to be overestimated unless phylogenetic comparative methods are employed^21,22^. Thus, when larger datasets are analysed with phylogenetic methods, the strength of catalytic trade-offs are substantially reduced^21,22^. Instead, phylogenetic constraints have had a larger impact on limiting enzyme adaptation compared to catalytic trade-offs^21,22^. These recent findings motivate a revaluation of the rubisco paradox, and an interrogation of whether rubisco is evolving for improved catalysis and CO_2_ assimilation in plants.

Here, we address these issues through a phylogenetic interrogation of the molecular and kinetic evolution of the Form I holoenzyme. We reveal that RbcL has evolved at a slower rate than >98% of all other gene/protein sequences across the tree of life. Through simultaneous evaluation of molecular and kinetic evolution of rubisco during the radiation of C_3_ angiosperms, we reveal that the enzyme has been continually evolving toward improved CO_2_/O_2_ specificity, carboxylase turnover rate, and carboxylation efficiency. Furthermore, we demonstrate that enhanced rubisco evolution is associated with enhanced rates of CO_2_ assimilation and higher photosynthetic nitrogen-use efficiencies. Thus, rubisco is not perfectly adapted, but is slowly evolving towards improved catalytic efficiency and CO_2_ assimilation.

## Results

### RbcL has evolved slower than RbcS and has experienced stronger purifying selection

Sequences encoding Form I rubisco were obtained from the National Center for Biotechnology Information (https://www.ncbi.nlm.nih.gov/). This dataset was filtered to retain sequences for a given species only if a full-length sequence for both *rbcL* and *rbcS* were present. Although *rbcL* exists as a single copy gene in all species, many species harbour multiple *rbcS* genes in their genomes. Thus, for each species a single *rbcL* sequence and all available *rbcS* sequences were taken forward. In total, this resulted in a set of 488 *rbcL*/RbcL and 1140 *rbcS*/RbcS sequences across 488 species (Supplemental File 1, Figure S1 and table S1).

In order to compare the rate at which the two rubisco subunits have evolved, species were partitioned into distinct taxonomic groups comprising the red algae (*Rhodophyta*; *n* = 201), the SAR supergroup (*Stramenopiles*, *Alveolates*, and *Rhizaria*; *n* = 129), the bacteria (*Bacteria*; *n* = 78), land plants (*Streptophyta*; *n* = 68) and green algae (*Chlorophyta*; *n* = 12) (Supplemental File 1, Figure S1 and table S1). Hereinafter, the total amount of molecular evolution of the nucleotide sequences (*rbcL* and *rbcS*) and the total amount of molecular evolution of the protein sequences (RbcL and RbcS) in a taxonomic group is referred to as “the extent of nucleotide evolution” and “the extent of protein evolution”, respectively. The term “the extent of molecular evolution” jointly refers to both.

Comparison of the two rubisco subunits revealed that the extent of molecular evolution in *rbcL*/RbcL is lower than that experienced by *rbcS*/RbcS (Figure 2A). This was not an artefact of the higher gene copy number of *rbcS,* as a 1,000 bootstrapped stratified sampling recovered the same result when only a single *rbcS/*RbcS sequence was randomly sampled per species (see Methods; Figure 2B). Therefore, *rbcL/*RbcL has explored less nucleotide and protein sequence space than *rbcS/*RbcS in the same sets of species over the same period of time (Figure 2C). Furthermore, *rbcL* also experienced fewer amino acid changes per change in nucleotide sequence compared to *rbcS* (Figure 2D), indicating a higher degree of purifying selection. Thus, *rbcL/*RbcL has evolved more slowly and has been subject to a higher degree of functional constraint on the encoded protein sequence than *rbcS/*RbcS.

**Figure 2.**
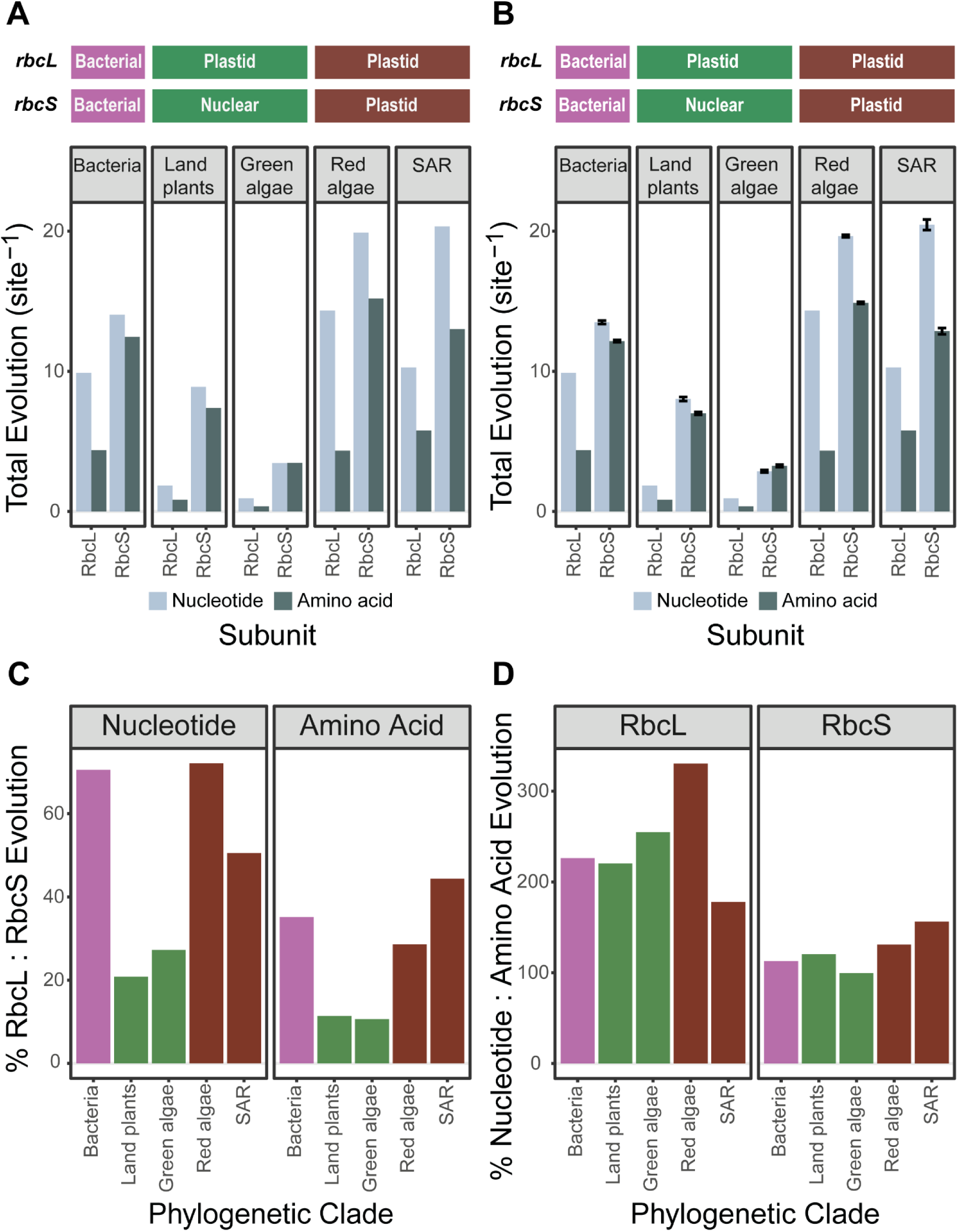
The extent of molecular evolution in rubisco during the radiation of each taxonomic group. **A)** Bar plot depicting the total amount of molecular evolution (substitutions per sequence site) in the nucleotide and protein sequences of the rubisco large (*rbcL*/RbcL) and small (*rbcS*/RbcS) subunit across taxonomic groups. The genome in which *rbcL* and *rbcS* genes reside within each group is indicated above the plot (bacterial, plastid, nuclear). **B)** As in (A) but using 1,000 bootstrapped stratified sampling of *rbcS*/RbcS per species to account for the higher copy number of this gene as compared to *rbcL*/RbcL in the dataset used for analysis (see Methods). Error bars represent ± 1 S.E of the mean. **C)** Bar plot depicting the percentage ratio (%) of nucleotide and amino acid evolution between rubisco subunits (*rbcL* to *rbcS* and RbcL to RbcS, respectively) in each taxonomic group. The colour of each bar is determined by the genome in which the *rbcL* and *rbcS* gene resides, following the colour scale in (A) and (B). **D)** Bar plot depicting the percentage ratio (%) of nucleotide to amino acid evolution in each rubisco subunit (*rbcL* to RbcL and *rbcS* to RbcS, respectively) in each taxonomic group. The colour of each bar is the same as described in (C).

### RbcL is one of the slowest evolving genes in the tree of life

To evaluate the rate of molecular evolution in context of all other genes in the species under consideration, the percentile rank of *rbcL*/RbcL and *rbcs*/Rbcs was evaluated for all genes in all species (see Methods). This revealed that 99.3% of all gene nucleotide sequences and 98.1% of all gene protein sequences evolved faster than *rbcL/*RbcL in the same sets of species over the same period of time (Figure 3A; Supplemental File 1, table S2). This held true even if *rbcL*/RbcL was only compared only to the subset of genes that encode enzymes, with 99.2% of enzyme nucleotide sequences and 98.3% of enzyme protein sequences evolving faster than *rbcL/*RbcL (Figure 3B; Supplemental File 1, table S3). Furthermore, in land plants *rbcL/*RbcL was also the slowest evolving component of the Calvin-Benson-Bassham cycle (Figure 3C; Supplemental File 1, table S4 and S5). This slow pace of evolution is not simply an artefact of being encoded in the plastid genome, as *rbcL/*RbcL was also one of the slowest evolving genes/proteins in bacteria which encode all of their genes in a single cytoplasmic genome. Thus, *rbcL/*RbcL is one of the slowest evolving genes/enzymes in all species in which it is found, irrespective of the taxonomic group or genome in which it is encoded.

**Figure 3.**
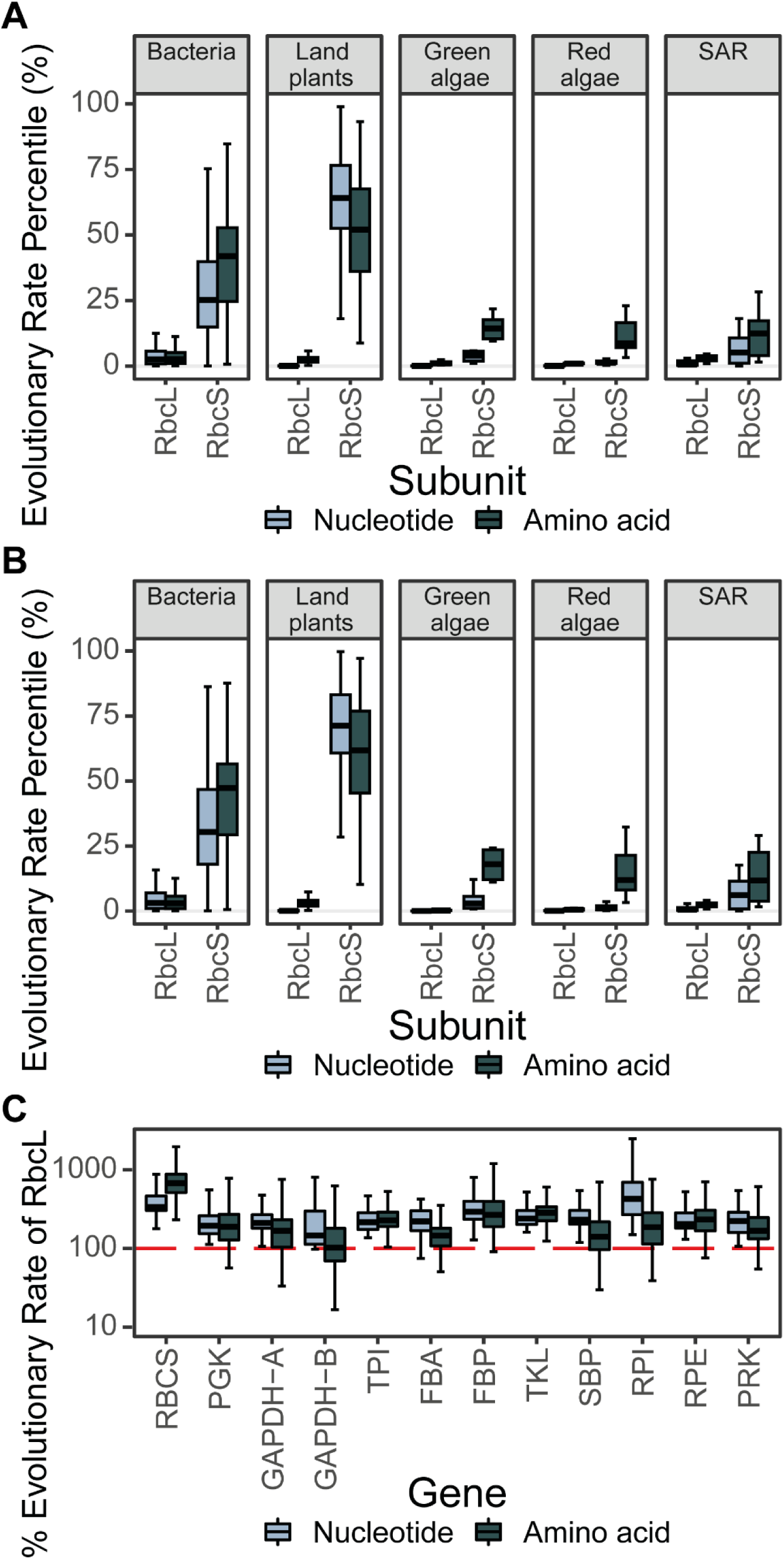
The extent of molecular evolution in rubisco in context other genes. **A)** Boxplot of the extent of molecular evolution (substitutions per sequence site) in the nucleotide and protein sequences of the rubisco large (*rbcL*/RbcL) and small (*rbcS*/RbcS) subunit expressed as a percentile (%) of that measured across all other genes and proteins, respectively. See also Supplemental File 1, table S2. **B)** As in (A) but calculating the percentile (%) extent of rubisco molecular evolution (substitutions per sequence site) relative to only the subset of genes and proteins in each species which encode enzymes. See also Supplemental File 1, table S3. **C)** Boxplot of the total amount of molecular evolution (substitutions per sequence site) in the nucleotide and protein sequences of each Calvin-Bensen-Bassham cycle enzyme expressed as a percentage (%) of that measured in *rbcL*/RbcL (100%; red horizontal line) across land plants. Phosphoglycerate kinase: PGK. Glyceraldehyde-3-phosphate dehydrogenase A/B subunit: GAPDH-A/GAPDH-B. Triose phosphate isomerase: TPI. Fructose-bisphosphate aldolase: FBA. Fructose-1,6-bisphosphatase: FBP. Transketolase: TKL. Sedoheptulose-bisphosphatase: SBP. Ribose 5-phosphate isomerase: RPI. Ribulose-p-3-epimerase: RPE. Phosphoribulokinase: PRK. See also Supplemental File 1, table S4 and table S5. The raw data for this figure can be found in Supplemental File 5.

In contrast to *rbcL*/RbcL, considerable variability in the extent of molecular evolution in the small subunit was observed both within and between taxonomic groups (Figure 3A; Supplemental File 1, table S2). Analogous results in each taxonomic group were recovered when this analysis was restricted to the subset of genes that encode enzymes (Figure 3B; Supplemental File 1, table S3). Moreover, in land plants *rbcS/*RbcS was the fastest evolving component of the Calvin-Benson-Bassham cycle (Figure 3C; Supplemental File 1, table S4 and S5). Thus, while the pace of molecular evolution in *rbcL/*RbcL is ubiquitously slow, the extent of molecular evolution of *rbcS/*RbcS is highly variable explaining the disparity in the rate of both subunits across the tree of life (Figure 2C; Supplemental File 1, table S6). A similar variable rate was also observed for rubisco’s ancillary chaperones (Supplemental File 1). Thus, the rate of molecular evolution of *rbcL*/RbcL is ubiquitously low, and lower than *rbcS*/RbcS or any associated chaperone.

### Rubisco is evolving for improved kinetic efficiency in plants

Although *rbcL* is among the slowest evolving genes on Earth, the analysis above demonstrates that it is not stagnant. This raises the question as to whether the sequence evolution is adaptive and is improving the catalysis of the enzyme. We hypothesised that if rubisco was undergoing directional selection for improved catalysis, then orthologs that have experienced the largest extent of molecular evolution would be the most efficient catalysts. To test this hypothesis, a dataset of kinetic measurements from C_3_ angiosperms^21,22,81^ was evaluated in the context of the molecular evolution of RbcL (Figure 4A,B). This analysis focused on RbcL as it is the primary determinant of kinetics^21–32^, and because sufficient sequence data for RbcS are unavailable. This revealed that the more RbcL has evolved from the most recent common ancestral sequence, the better its CO_2_/O_2_ specificity (*S*_C/O_; 10.1% variance explained), CO_2_ turnover rate (*k*_catC_; 4.6% variance explained) and carboxylation efficiency (*k*_catC_/*K*_C_; 3.8% variance explained) (Figure 4B). This result is not an artefact caused by potential systematic methodological biases associated with species sampling or potential uncertainties or errors in the underlying phylogenetic tree (See Methods, Supplemental File 1). Thus, rubisco has been adaptively evolving for improved *S*_C/O_, *k*_catC_, and *k*_catC_/*K*_C_ during the radiation of the angiosperms.

**Figure 4.**
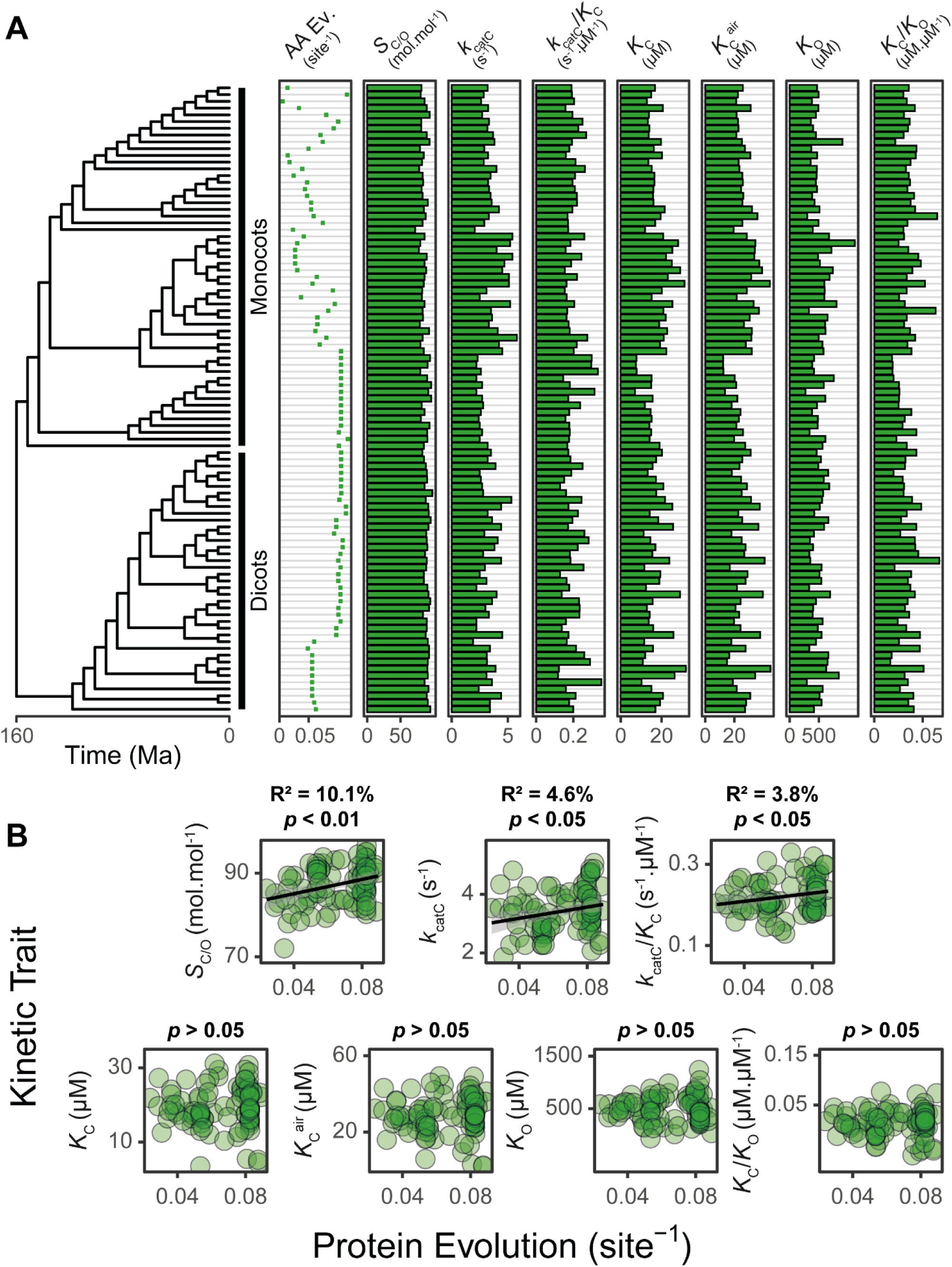
The relationship between rubisco molecular and kinetic evolution in C_3_ angiosperms. **A)** The relationship between RbcL evolution and its corresponding kinetic trait values. AA Ev.: The extent of RbcL amino acid evolution that has occurred since the last common ancestor at the root of the angiosperm phylogeny. S_C/O_: specificity. *k*_catC_: carboxylase turnover per site. *k*_catC_/*K*_C_: carboxylation efficiency. *K*_C_: the Michaelis constant for CO_2_. *K*_Cair_: the inferred Michaelis constant for CO_2_ in 20.95% O_2_. *K*_O_: the Michaelis constant for O_2_. *K*_C_/*K*_O_: the ratio of the Michaelis constant for CO_2_ compared to O_2_. **B)** The relationship between the extent of RbcL protein evolution (substitutions per sequence site) and each rubisco kinetic trait in (A) as assessed using least squares regression models. The raw data can be found in Supplemental File 7.

Given that the origin of the angiosperms is estimated to have occurred 160 million years ago^84^ (Figure 4A), it is possible to put the above kinetic change in context of both molecular sequence changes and evolutionary time (table 1). As the large subunit acquired one nucleotide substitution every 0.9 million years and one amino acid substitution every 7.2 million years (Supplemental File 1, figure S2), each amino acid substitution resulted in an average increase in *S*_C/O_ by 2.7×10^-1^ mol.mol^-1^, in *k*_catC_ by 3.6×10^-2^ s^-1^, and in *k*_catC_/*K*_C_ by 1.8×10^-3^ s^-1^ µM^-1^. This is equivalent to a relative improvement of 0.3% (*S*_C/O_), 1.4% (*k*_catC_), and 1.1% (*k*_catC_/*K*_C_) per amino acid substitution, and a relative improvement of 0.05% (*S*_C/O_), 0.2% (*k*_catC_), and 0.2% (*k*_catC_/*K*_C_) per million years. Thus, there has been continual improvement in rubisco kinetics during the radiation of the angiosperms at a rate that is dependent on the extent of its molecular sequence change.

**Table 1.**
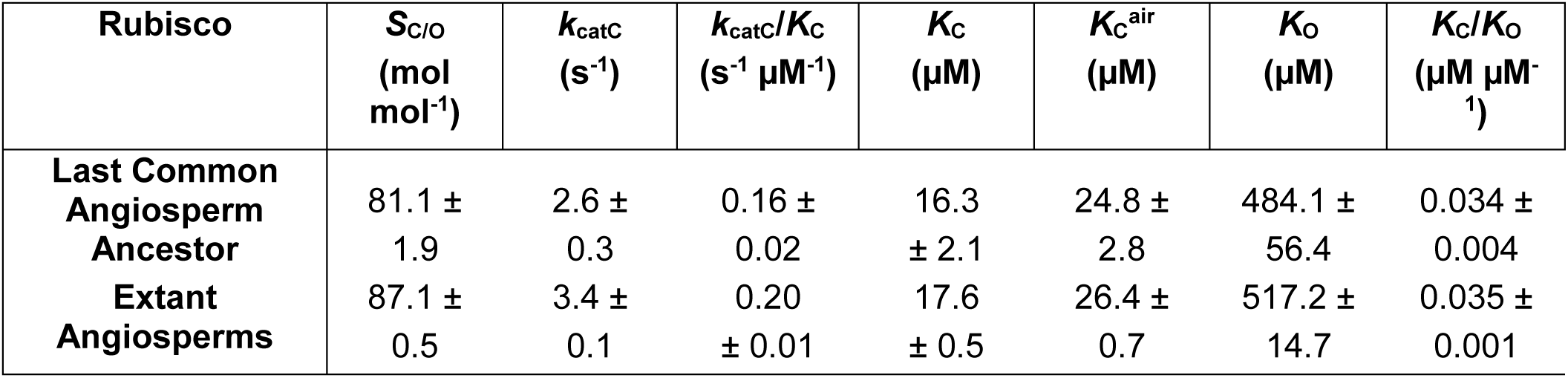
Rubisco kinetics in extinct and extant angiosperms. Kinetic trait values for the last common ancestor of the angiosperms were computed based on the estimated y intercept (mean ± 1 S.E.) of the linear regression analysis performed between the extent of RbcL protein evolution and each rubisco kinetic trait in Figure 4B. Mean values of rubisco kinetic traits and associated variation (± 1 S.E.) in extant C_3_ species are shown for comparison. The raw data set used can be found in Supplemental File 7.

| Rubisco | $S_{C/O}$<br>(mol<br>mol<sup>-1</sup>) | $k_{catC}$<br>(s<sup>-1</sup>) | $k_{catC}/K_C$<br>(s<sup>-1</sup> μM<sup>-1</sup>) | $K_C$<br>(μM) | $K_C^{air}$<br>(μM) | $K_O$<br>(μM) | $K_C/K_O$<br>(μM μM<sup>-1</sup>) |
| --- | --- | --- | --- | --- | --- | --- | --- |
| **Last Common**<br>**Angiosperm**<br>**Ancestor** | 81.1 ±<br>1.9 | 2.6 ±<br>0.3 | 0.16 ±<br>0.02 | 16.3<br>± 2.1 | 24.8 ±<br>2.8 | 484.1 ±<br>56.4 | 0.034 ±<br>0.004 |
| **Extant**<br>**Angiosperms** | 87.1 ±<br>0.5 | 3.4 ±<br>0.1 | 0.20<br>± 0.01 | 17.6<br>± 0.5 | 26.4 ±<br>0.7 | 517.2 ±<br>14.7 | 0.035 ±<br>0.001 |

### Rubisco is evolving for improved leaf-level CO2 assimilation

Given that rubisco is evolving to become a better catalyst, we hypothesised that this adaptation would also drive adaptation in the rate of leaf-level CO_2_ assimilation. To test this hypothesis we analysed a large dataset of photosynthetic measurements from C_3_ angiosperms^85^ in the context of the extent of their RbcL evolution (Figure 5A-C). This revealed that the rate of leaf-level CO_2_ assimilation was also dependent on the extent of molecular sequence change in rubisco such that that C_3_ angiosperms with more evolved rubisco also higher rates of CO_2_ assimilation (*A*_mass_; 19.2% variance explained, Figure 5B). This is not a consequence of increased nitrogen investment in the leaf, as the association between rubisco evolution and increased CO_2_ assimilation is strengthened when measurements are controlled for leaf nitrogen content (*PNUE_mass_,* 22.1% variance explained, Figure 5B). Analogous results were obtained when measurements of CO_2_ assimilation were evaluated on a leaf area basis (Figure 5C). This result is most parsimoniously explained by directional selection towards enhanced leaf-level CO_2_ assimilation driven by the kinetic adaptation described above. Thus, the adaptive evolution of rubisco during the radiation of the angiosperms has resulted in the improvement in leaf-level CO_2_ assimilation.

**Figure 5.**
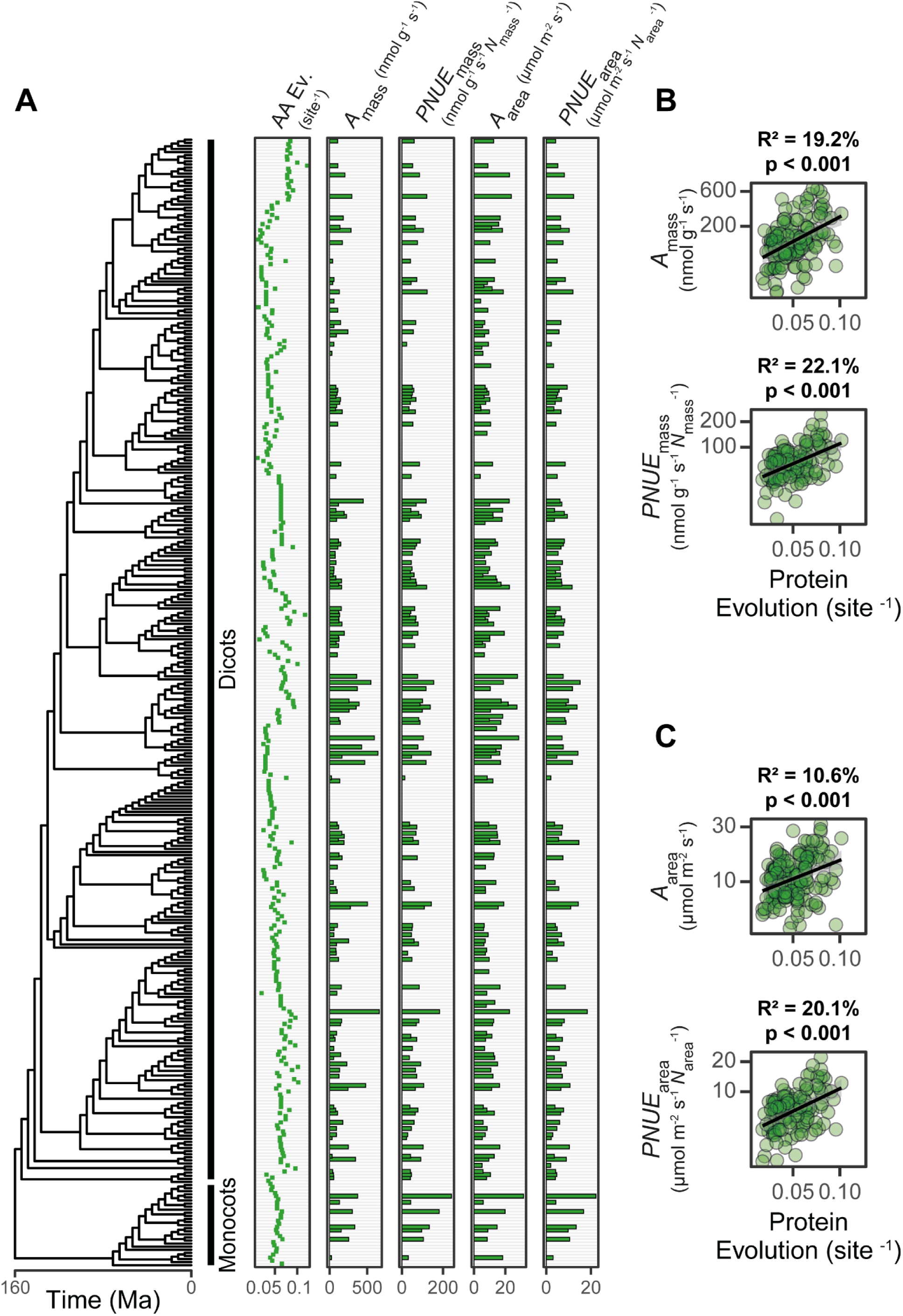
The relationship between rubisco molecular evolution and CO_2_ assimilation in C_3_ angiosperms. **A)** The relationship between the extent of RbcL evolution and leaf level CO_2_ assimilation. AA Ev.: The extent of RbcL amino acid evolution that has occurred since the most recent common ancestor at the root of the angiosperm phylogeny. *A*_mass_: Photosynthetic rate per unit leaf mass. *PNUE*_mass_: Photosynthetic nitrogen use efficiency rate per unit leaf mass per unit leaf nitrogen content (*N*_mass_; % N). *A*_area_: Photosynthetic rate per unit leaf area. *PNUE*_area_: Photosynthetic nitrogen use efficiency rate per unit leaf area expressed per unit leaf area nitrogen content (*N*_area_; g m^-2^ N). **B)** The relationship between the extent of RbcL protein evolution (substitutions per sequence site) and each photosynthetic trait in (A) evaluated on a mass-basis (*A*_mass_, *PNUE*_mass_) as assessed using least squares regression models. **C)** As in (B) but for each photosynthetic trait evaluated on an area-basis (*A*_area_, *PNUE*_area_). The raw data can be found in Supplemental File 9.

## Discussion

Rubisco is the primary entry point for carbon into the biosphere, responsible for fixing 250 billion tons of CO_2_ annually^18^. Despite this immense throughput, the enzyme is a surprisingly inefficient catalyst with a modest carboxylase turnover rate of <12 s^-1^ ^60^ and a competing oxygenase activity that results in the loss of fixed carbon^63,64,86^. This discord presents an evolutionary paradox that has attracted significant attention^15,21,22,63,67,74–78^, with the prevailing assumption being that rubisco is evolutionarily stagnant. Here we demonstrate that the enzyme is not stagnant, but that it is encoded by one of the slowest evolving genes on Earth. Despite this, we demonstrate that rubisco has been evolving for higher CO_2_/O_2_ specificity (*S*_C/O_), faster carboxylase turnover rates (*k*_catC_), and improved carboxylation efficiencies (*k*_catC_/*K*_C_) in angiosperms. Moreover, we demonstrate that plants with more evolved rubisco exhibit higher leaf-level CO_2_ assimilation and enhanced photosynthetic nitrogen-use efficiencies. Thus, rubisco has been continually evolving towards improved catalytic efficiency and CO_2_ assimilation during the radiation of the angiosperms.

A slow rate of molecular evolution in *rbcL* has long been assumed and has underpinned the use of this gene for systematics and phylogenetics^87–89^. However, to our knowledge there has been no contextualised measurement of the rate of *rbcL* evolution across the tree of life. The analysis presented here addresses this gap by revealing that across the tree of life, *rbcL*/RbcL has experienced a lower extent of molecular evolution than 99% of all gene nucleotide sequences and 98% of all gene protein sequences. It is interesting to note here that this is not due to the presence of *rbcL* in the chloroplast genome, as *rbcL* is also one of the slowest evolving sequences in bacteria which lack organellar genomes. Thus, RbcL is universally one of the slowest evolving sequences on Earth, irrespective of the taxon or genome in which it resides.

Although dissecting the factors which constrain the rate of *rbcL* evolution is beyond the scope of the current study, the slow pace of *rbcL* molecular evolution is most likely a consequence of several synergistic factors^90^ including constraints imposed by expression^91–95^, selection to preserve protein function^96–100^, and the requirements for protein-protein interactions *in vivo*^101–104^. These factors would be particularly pertinent for rubisco given that it is the most abundant protein in organisms in which it is found^68,70^, it is subject to catalytic trade-offs^21,22,80,81^ and molecular activity-stability trade-offs^105–108^, and given that it relies on multiple interacting partners and chaperones for folding, assembly and metabolic regulation^68,109^. Thus, a perfect storm of features exist which could limit the molecular evolution of *rbcL* and thereby cause it to be one of the slowest evolving genes on Earth. Further work to elucidate the exact contribution of each of these biological determinants on rubisco’s rate of molecular evolution is warranted, building upon the work here and previous investigations^21,22,110^.

Our integrated analysis of rubisco evolution revealed a continual improvement in *S*_C/O_, *k*_catC_ and *k*_catC_/*K*_C_ during the radiation of C_3_ angiosperms. Thus, although rubisco is slowly-evolving, sequence changes have enhanced the catalytic properties of the enzyme. In the context of the C_3_ leaf, such directional selection towards improved *S*_C/O_ is consistent with adaptation to maintain adequate carbon assimilation in response to declining CO_2_ and increasing O_2_ (Figure 1). This evolutionary strategy has been proposed previously^80^, and is suggested to apply broadly across photoautotrophs lacking a CO_2_ concentrating mechanism^82,111^. In addition to adaptation for higher *S*_C/O_, we also discover simultaneous improvement in *k*_catC_ and *k*_catC_/*K*_C_ without antagonism in any other kinetic trait. These results are also consistent with the inferior *S*_C/O_ and *k*_catC_/*K*_C_ reported for extinct rubisco resurrected at the dawn of the Form IB^6^ and Form 1^34^ lineages. It is noteworthy that on first appearances, all of these studies seem to contradict an analysis within the *Solanaceae* in which resurrected ancestral rubisco variants exhibited superior *k*_catC_ and *k*_catC_/*K*_C_ values. However, in this instance the kinetic differences were proposed to be driven by sequence changes in RbcS^112^, and therefore do not contradict the analysis of RbcL presented here or in other studies^6,34^. Thus, sequence change in RbcL during the radiation of angiosperms has driven the continual improvement of the enzyme in the presence of a declining atmospheric CO_2_:O_2_ concentration.

The ‘FvCB model’ of photosynthesis^113^, as well as a suite of other experimental^114–121^ and computational^122,123^ studies all demonstrate that rubisco is a major rate-limiting factor for CO_2_ assimilation under ambient steady-state conditions. The findings presented here link these mechanistic studies with evolutionary biology, and reveal that rubisco has experienced directional selection to improve kinetic efficiency and CO_2_ assimilation. Ultimately, this changes our understanding of the rubisco paradox. Rubisco is not locked in evolutionary stasis, but is instead slowly evolving towards improved CO_2_ assimilation. This discovery has significant implications for our understanding of the past, present, and future potential of rubisco in natural and engineered contexts.

## Materials and Methods

### Rubisco sequence data

All publicly available coding sequences of the rubisco large (*rbcL*) and small (*rbcS*) subunit genes in the National Center for Biotechnology Information (NCBI) database (https://www.ncbi.nlm.nih.gov/) as of July 2020 were downloaded. Manual inspection of nucleotide and translated protein sequences was performed to remove any partial, chimeric, or erroneously annotated sequences. In addition, this dataset was further restricted to include only those species which possess a Form I rubisco and for which both a full-length *rbcL* and *rbcS* gene sequence could be obtained. Given that *rbcL* exists as a single copy gene in all species, only one *rbcL* sequence per species was retained for downstream analysis. In contrast, all possible full-length *rbcS* sequences were taken forward to account for the fact that *rbcS* is multicopy in some genomes.

Translated *rbcL* and *rbcS* protein sequences were aligned using the MAFFT L-INS-i algorithm^124^. The corresponding codon alignments of the nucleotide sequences were generated by threading the nucleotide sequences through the aligned protein sequences that they encode using PAL2NAL software^125^. Multiple sequence alignments were trimmed to remove non-aligned codon or residue positions such that only ungapped columns remained. During this process, the putative transit peptide of the *rbcS*/RbcS sequences in taxa in which this gene is encoded by the nuclear genome was computationally cleaved. Following these data processing steps, alignments were partitioned depending on species membership to either the bacteria (*Bacteria*; *n* = 78), land plants (*Streptophyta*; *n* = 68), green algae (*Chlorophyta*; *n* = 12), red algae (*Rhodophyta*; *n* = 201) or the SAR supergroup (*Stramenopiles*, *Alveolates*, and *Rhizaria; n* = 129) by use of the NCBI taxonomy browser (https://www.ncbi.nlm.nih.gov/Taxonomy/TaxIdentifier/tax_identifier.cgi). Any sequences belonging to species in either the *Haptophyta*, *Cryptophyta*, *Glaucocystophyta* or *Excavata* taxonomic groups were excluded from the dataset at this point due to insufficient data availability. In total, this resulted in a combined set of 488 *rbcL*/RbcL and 1140 *rbcS*/RbcS gene and protein sequences across 488 species spanning 5 taxonomic groups (Supplemental File 1, Figure S1 and table S1). The complete set of raw *rbcL*/RbcL and *rbcS*/RbcS sequences, as well as the complete set of aligned and trimmed *rbcL*/RbcL and *rbcS*/RbcS sequences can be found in Supplemental File 2.

### Rubisco phylogenetic tree inference

Maximum-likelihood *rbcL*/RbcL and *rbcS*/RbcS phylogenetic gene trees were inferred across all sequences within a taxonomic group by IQ-TREE^126^ using the ultrafast bootstrapping method with 1,000 replicates and the Shimodaira–Hasegawa approximate–likelihood ratio branch test. The best fitting models of nucleotide (SYM+R8) and amino acid (LG+R5) sequence evolution were respectively determined as those which exhibit the lowest combined Bayesian information criterion rank score across the complete sets of both rubisco large and small subunit sequences^126^. Across all taxonomic groups, the models of nucleotide and amino acid sequence evolution were held constant between the gene trees for *rbcL* and *rbcS*, and RbcL and RbcS, respectively, such that branch lengths are comparable across both subunits. The complete set of these *rbcL*/RbcL and *rbcS*/RbcS phylogenetic gene trees used as the basis of the analysis herein can be found in Supplemental File 3.

### Stratified sampling of rbcS sequences and phylogenetic tree inference

To account for potential biases in our analysis caused by some species exhibiting multiple copies of *rbcS*, random stratified sampling of the non-gapped *rbcS*/RbcS sequence alignments was conducted by species using 1,000 replicates with replacement. This process resulted in the generation of 1,000 unique *rbcS*/RbcS alignments for each taxonomic group, whereby each of these respective alignments contain only a single randomly selected copy of the *rbcS*/RbcS per species. In turn, each of these alignments were subject to data processing and phylogenetic tree inference using IQ-TREE^126^ following the method described above.

### Quantification of the total extent of nucleotide and protein molecular evolution in rbcL/RbcL and rbcS/RbcS

The extent of molecular evolution in both rubisco subunits was assessed across all species in a given taxonomic group as the total length (sequence substitutions per aligned sequence site) of the phylogenetic tree describing the evolutionary history of each respective gene. For this purpose, tree length was calculated as the combined sum of branch lengths leading from the root at the last common ancestor of the tree to the set of sequences at the terminal nodes. In this way, using the trees inferred across the complete cohort of *rbcL*/RbcL and *rbcS*/RbcS sequences in each taxonomic group, it was possible to capture all nucleotide and amino acid evolution which has arisen in each subunit since the most recent common ancestor of all sampled species in the group. An identical analysis was also performed for each *rbcS*/RbcS tree generated by stratified sampling, with mean and standard errors of estimates being calculated in this case across the 1,000 unique bootstrap replicate trees.

### Genomes and gene models

Complete sets of representative gene models for as many species in the rubisco sequence dataset as possible were acquired from either NCBI (https://www.ncbi.nlm.nih.gov/) or Phytozome V13^127^.

Where more than one such gene model resource was available for a given species, the most recent assembly version was chosen. In this way, complete sets of representative gene models were acquired for a total of 32 of the bacteria species, 27 of the land plant species, 8 of the SAR species, 6 of the red algae species and 4 of the green algae species analysed in the present study, respectively (Supplemental File 1, table S1 and S7).

Predicted gene model sets were filtered to remove sequences with internal in-frame stop codons. Gene model sets were also filtered to keep only the longest gene model variant per gene. Moreover, owing to a lack of data availability of publicly available chloroplast or mitochondrial genomes for the eukaryotic species in the present analysis, and as organellar genomes contain fewer than 1% of genes encoded in the corresponding nuclear genome, only gene sequences encoded by the nuclear genomes of species in the land plant, green algae, red algae and SAR taxonomic groups were taken forward for analysis. Finally, after the above quality control checks were completed, a corresponding proteome was generated from each species gene model set by *in silico* translation of the respective coding sequences.

### Orthogroup classification and phylogenetic tree inference

The complete set of translated proteomes for species in each respective taxonomic group were subject to orthogroup inference using OrthoFinder V2.5.2^128,129^ software run with default settings and with the DIAMOND ultra-sensitive mode^130^. Protein sequences within each orthogroup were aligned using the MAFFT L-INS-I algorithm with 1,000 cycles of iterative refinement^124^. The corresponding codon alignments of the nucleotide sequences were generated by threading the nucleotide sequences through the aligned protein sequences that they encode using PAL2NAL software^125^. Alignments were trimmed to remove positions which contain gap characters. Sequences that were <50% of the median length of the cohort of all other sequences in the given orthogroup were excluded to avoid analysis of partial or truncated genes that could influence downstream analysis. All nucleotide and protein multiple sequence alignments which satisfied the above criteria and which possessed >50 ubiquitously aligned codon or amino acid positions were subject to bootstrapped maximum likelihood phylogenetic tree inference using IQ-TREE^126^ following the exact method and evolutionary substitution models described above. In total, this resulted in a combined set of 16,631 orthogroup phylogenies comprising 5,126,017 ortholog pairwise comparisons across 351 species pairwise comparisons for the land plant clade, 6,953 orthogroup phylogenies comprising 153,288 ortholog pairwise comparisons across 28 species pairwise comparisons for the SAR clade, 5,422 orthogroup phylogenies comprising 642,057 ortholog pairwise comparisons across 496 species pairwise comparisons for the bacteria clade, 4,269 orthogroup phylogenies comprising 31,133 ortholog pairwise comparisons across 6 species pairwise comparisons for the green algae clade and 3,966 orthogroup phylogenies comprising 54,091 ortholog pairwise comparisons across 15 species pairwise comparisons for the red algae clade, from which to base the analyses herein. A further breakdown of these metrics for each species comparison can be found in Supplemental File 4.

### Characterization of the set of enzymatic gene and protein sequences within orthogroups

The set of all genes within each species proteome that encode enzymes was determined using the DeepEC^131^ deep learning-based classifier algorithm. For this purpose, enzymes were defined as those protein sequences that could be assigned at least a partial enzyme commission (EC) number (i.e., at minimum, a single digit EC top-level code). On average 42.2% of all genes in the analysis encoded enzymes. A detailed breakdown of the metrics of enzyme ortholog pairwise comparisons for each species comparison can be found in Supplemental File 4.

### Quantification of percentiles of the rate of molecular evolution

To evaluate the extent of molecular evolution in rubisco in the context of all other genes, only species in the rubisco sequence dataset possessing a publicly available whole-genome gene assembly were considered. Across each pairwise combination of species in a given taxonomic group which satisfied this criteria, the extent of *rbcL*/RbcL and *rbcS*/RbcS molecular evolution since the time point of species divergence was measured by computing the sum of branch lengths (sequence substitutions per aligned sequence site) separating these respective sequences in the rubisco phylogenetic trees previously inferred. Following this, the extent of molecular evolution separating all other pairs of orthologous (but not paralogous) gene and protein sequences for that given species pair was measured across all inferred orthogroup phylogenies, and the percentile rank rate of rubisco nucleotide or protein evolution was computed relative to the cohort of these measurements. To assess the extent of rubisco molecular evolution in context all other enzymes, the exact same steps were followed but only the subset of genes and proteins predicted to encode enzymes were included. In both of these analyses, a minimum threshold of 100 measurements for orthologous genes and protein sequences was ensured per species pair. In cases where multiple percentiles are calculated for a rubisco subunit in a given species pair (due to gene duplications in the *rbcS* of some species, or due to a single species gene assembly matching multiple sub-species in the rubisco sequence dataset) the mean percentile was taken. The full set of data generated from these analyses quantifying the relative percentile extent of rubisco molecular evolution to all other genes and proteins, and to all other enzyme-encoding genes and proteins can be found in raw and processed forms in Supplemental File 5.

### Identification and classification Calvin-Benson-Bassham cycle enzyme isoforms in land plants

The set of genes which encode Calvin-Benson-Bassham cycle enzymes was first resolved in the model plant species *Arabidopsis thaliana*. To achieve this, the complete gene families to which each Calvin-Benson-Bassham cycle enzyme in *A. thaliana* belongs was determined based on available data in The Arabidopsis Information Resource (TAIR) database (http://arabidopsis.org)^132,133^. Following this, the photosynthetic isoforms in these gene families which are active in the Calvin-Benson-Bassham cycle in the chloroplast stroma were then identified based on several lines of evidence. 1) A high protein abundance based on whole-organism integrated protein abundance data obtained from the Protein Abundance Database (https://pax-db.org/) dataset 3702/323. 2) Leaf mRNA expression based on tissue-specific RNA sequencing data obtained from both the Arabidopsis eFP Browser V2.0 (http://bar.utoronto.ca/efp2/Arabidopsis/Arabidopsis_eFPBrowser2.html) and the EMBL-EBI (https://www.ebi.ac.uk/) dataset E-GEOD-53197. 3) Chloroplast-targeted protein subcellular localisation as predicted using both TargetP V2.0^134,135^ and Predotar V1.04^136^. 4) Gene orthology as inferred from trees generated for each Calvin-Benson-Bassham cycle gene family using IQ-TREE^126^. The resulting set of photosynthetic isoforms encoding each Calvin-Benson-Bassham cycle enzyme in *A. thaliana* inferred from this multi-faceted analytical pipeline can be found in Supplemental File 1, table S8.

The set of genes which encode the photosynthetic isoforms of Calvin Bensen Bassham cycle enzymes in all other 26 land plant species (apart from *A. thaliana*) for which genome sequence data was available in this study were then determined by orthology using data from the orthogroup inference analysis performed above. Each group of orthologous protein sequences determined to encode a given Calvin-Benson-Bassham cycle enzyme across all species were aligned using the MAFFT L-INS-i algorithm^124^, and corresponding nucleotide coding sequence alignments were generated using PAL2NAL^125^. Multiple sequence alignments were subject to the same data filtering and quality control criteria previously described to remove partial or incomplete sequences and subsequently delete any column positions which contain gaps. Finally, bootstrapped maximum likelihood phylogenetic trees were inferred by IQ-TREE^126^ following the method outlined above. A similar analysis of Calvin-Bensen-Bassham cycle enzymes in other taxonomic groups was not able to be performed owing to a lack of the required data described here to determine the photosynthetic gene isoforms in these species.

### Quantification of the relative extent of rbcL/RbcL and rbcS/RbcS molecular evolution relative to all Calvin-Benson-Bassham cycle enzymes in land plants

To determine the extent of molecular evolution between orthologous Calvin-Benson-Bassham cycle isoforms compared to Form I rubisco, the extent of molecular evolution measured between all other Calvin-Benson-Bassham cycle gene and protein orthologous sequences for each species pair were expressed as a percentage ratio of that measured in the corresponding *rbcL*/RbcL sequence. In cases where multiple percentage ratios are calculated for a given Calvin-Bensen-Bassham component in a given species pair (due to gene duplications, or due to a single species gene assembly matching multiple sub-species in the rubisco sequence dataset) the mean value was taken. The full set of data generated from this analysis can be found in raw and processed forms in Supplemental File 5.

### Quantification of the percentile extent of rubisco chaperone molecular evolution within each taxonomic group

To evaluate the percentile rate of molecular evolution in the known chaperones of Form I rubisco in the context of all other genes in each taxonomic group, the exact same method was followed as above for *rbcL*/RbcL and *rbcS*/RbcS though the subject of the analysis was respectively altered. Here, for this investigation, the putative set of genes which encode each Form I ancillary chaperone involved in holoenzyme metabolic regulation (RUBISCO ACTIVASE (Rca)) and in holoenzyme folding and assembly (BUNDLE SHEATH DEFECTIVE 2 (BSD2), CHAPERONIN 10 (Cpn10), CHAPERONIN 20 (Cpn20), CHAPERONIN-60 (Cpn60), RBCX (RbcX), RUBISCO ACCUMULATION FACTOR 1 (Raf1), RUBISCO ASSEMBLY FACTOR 2 (Raf2)) were first resolved in the model plant species *A. thaliana*. This was achieved using a previously published dataset^137^ supplemented by information available in the TAIR database (http://arabidopsis.org)^132,133^. The resulting set of *Arabidopsis* chaperone genes thus identified can be found in Supplemental File 1, table S9. Following this step, the corresponding set of genes encoding rubisco chaperones in all other species for which a complete gene assembly could be obtained were inferred using data from a separate OrthoFinder run performed with identical settings as previously described, but based on the translated proteomes of all organisms across all taxonomic groups.

After the cohort of Form I rubisco chaperone genes were identified in all species, the percentile rates of nucleotide and protein evolution in these genes were calculated between each pairwise combination of species relative to all other pairs of orthologous sequences using the identical measurements previously generated from the analysis of *rbcL*/RbcL and *rbcS*/RbcS above. In this way, analysis of some chaperones were omitted in certain taxonomic groups owing to the data quality and filtering steps that were previously performed as described above. In cases where multiple percentiles are calculated for a chaperone in a given species pair (due to gene duplications, or due to a single species gene assembly matching multiple sub-species in the rubisco sequence dataset) the mean percentile was taken as above. The full set of data generated from this analysis can be found in raw and processed forms in Supplemental File 5. A combined dataset including the relative percentile extent of evolution in both rubisco subunits and all rubisco chaperones for each unique pairwise species comparison can be found in Supplemental File 6.

### Integrated analysis of rubisco molecular and kinetic evolution

To interrogate the relationship between the molecular and kinetic evolution of extant Form I rubisco, a dataset of rubisco kinetic traits was downloaded from^21,22^, as modified from that originally compiled by Flamholz and colleagues^81^. For the purpose of this study, only species in this dataset with a complete set of experimentally determined measurements of rubisco specificity (*S*_C/O_) for CO_2_ relative to O_2_ (i.e., the overall carboxylation/oxygenation ratio of rubisco under defined concentrations of CO_2_ and O_2_ gases), maximum carboxylase turnover rate per active site (*k*_catC_), and the respective Michaelis constant (i.e., the substrate concentration at half-saturated catalysed rate) for both CO_2_ (*K*_C_) and O_2_ (*K*_O_) substrates were selected. For each of the 137 species which satisfied this criteria (all of which were angiosperm land plants), an estimate of the Michaelis constant for CO_2_ in 20.95% O_2_ air (*K*_Cair_) was also available^21,22^. In addition, the ratio of the Michaelis constant for CO_2_ relative to O_2_ (*K*_C_/*K*_O_), as well as carboxylation efficiency, defined as the ratio of the maximum carboxylase turnover to the Michaelis constant for CO_2_ (*k*_catC_/*K*_C_), were also inferred. Measurements of the Michaelis constant for RuBP (*K*_RuBP_) were not considered owing to a limited sample size (*n* = 19). All *Limonium* species in the dataset were also ignored on the basis that trait values obtained across different studies have been deemed to not be consistent^24,138^. In total, this left a dataset of rubisco kinetic trait measurements for 123 angiosperms. Of these, only the subset of 93 species which perform C_3_ photosynthesis were considered for the purpose of the integrated molecular and kinetic evolution analysis herein. This is because of both a limited sample size of C_3_-C_4_ (*n* = 6), C_4_-like species (*n* = 3) and C_4_ species (*n* = 21) in the kinetic dataset, and given that transition toward C_4_ photosynthesis is associated with a change in rubisco kinetic evolution^21,22^ that would confound the directional selection analysis being conducted.

Coding sequences of the *rbcL* gene were obtained from^22^ for each species in the kinetic dataset. In order to facilitate more accurate downstream phylogenetic tree inference across these sequences and to minimize the impact of long-branch effects^139^, the complete set of publicly available *rbcL* coding sequences in land plants were also acquired in parallel from NCBI (https://www.ncbi.nlm.nih.gov/) using the query term “rbcL[Gene Name] AND “plants”[porgn:_txid3193]”. These sequences thus obtained were subject to the exact same data processing steps to remove ambiguous, partial or chimeric sequences as performed previously for the *rbcL* sequences of species in the rubisco kinetic dataset^22^. In total, this step resulted in an additional set of 29,218 full-length *rbcL* coding sequences to aid downstream phylogenetic inference. Protein sequences were inferred from each *rbcL* coding sequences via *in silico* translation. Next, the complete set of translated RbcL sequences (including the set of sequences from angiosperms in the rubisco kinetic dataset, as well as the set of all publicly available sequences for land plants) were respectively aligned using MAFFT L-INS-I^124^, and a corresponding *rbcL* coding sequence alignment was generated using PAL2NAL software^125^. The resulting multiple sequence alignments were trimmed to remove non-aligned residue positions and bootstrapped phylogenetic trees were inferred using IQ-TREE^126^ following the exact method described above and using the best-fit models of nucleotide and protein sequence evolution previously inferred. To facilitate downstream analysis, the *rbcL* and RbcL gene trees were subsequently modified to keep only internal and terminal branches leading to the set of species in the rubisco kinetic dataset, with pruned trees manually rooted in Dendroscope^140^.

To compute the relative extent of protein evolution which has occurred in each angiosperm in the kinetic dataset, the summed branch length (sequence substitutions per aligned sequence site) leading from the last common ancestor at the root of this clade to each respective terminal node in the RbcL phylogeny generated above were measured. The kinetic trait values and extent of molecular evolution for all C_3_ angiosperm rubisco can be found in Supplemental File 7. The predicted kinetic trait values at the last common ancestor at the base of the angiosperm clade were inferred from the estimated y intercept values from these regression models and can be found in table 1. The *rbcL*/RbcL phylogenetic gene trees used as the basis of this analysis, including the trees inferred across the full set of sequences, as well as the pruned versions of these trees containing only the subset of C_3_ species in the kinetic dataset, can be found in Supplemental File 8.

To assess the robustness of the above integrated molecular and kinetic investigation of rubisco evolution, the same analysis was repeated but including only the minimal subset of species in the kinetic dataset which captured the majority of phylogenetic diversity across all sampled species (see below), so as to control for biases associated with species sampling and overrepresentation of certain groups (see Supplemental File 1). An identical analysis was also performed using the complete set of species in the kinetic dataset but based on analogous trees generated following the exact same method as above but based on alternate best-fitting models of sequence evolution inferred for the specific alignment, so as to control for potential artefacts associated with errors or uncertainties in phylogenetic tree inference (Supplemental File 1). As the results of these supplementary analyses were identical to that generated from our original analysis, our conclusions were demonstrated to be valid and robust and not an artefact caused by either systematic biases in species sampling or by errors in phylogenetic reconstruction.

### Accounting for potential species sampling error

To identify a minimal subset of species which capture all of the phylogenetic diversity (PD) contributed to by the complete set of 93 C_3_ species in the rubisco kinetic dataset, the Phylogenetic Diversity Analyzer V1.0.3 software^142^ was employed using the ‘greedy’ algorithm. Specifically, the unrooted RbcL phylogenetic tree of the 93 C_3_ species was subject to systematic interrogation to identify the optimal combination of species at each iterative tree size (*n* = 2 – 93 species) which maximizes the PD score. For the purpose of this method, PD is defined as the total tree length (i.e., the combined sum of all internal and terminal branch lengths) of the pruned phylogeny comprising the selected subset of sampled species. Based on the results of this analysis, it was observed that phylogenetic diversity saturated at a threshold of 50 species (accounting for 54.8% of the total 93 C_3_ species in the kinetic dataset) (Supplemental File 1). The optimal composition of species at this respective threshold included 31 dicotyledonous individuals and 19 monocotyledonous individuals, and are listed in Supplemental File 1, table S10. This set of 50 species were used to assess the robustness of the molecular and kinetic analysis of rubisco to potential artefacts associated with biases in species sampling.

### Integrated analysis of rubisco molecular evolution and CO2 assimilation

To investigate the relationship between rubisco molecular evolution and whole-plant photosynthetic performance, a comprehensive meta-dataset of photosynthetic measurements from species spanning the whole land plant phylogeny was provided by Gago and colleagues^85^. This dataset contained measurements of light-saturated net photosynthetic rates expressed both per unit leaf mass (*A*_mass_) and per unit leaf area (*A*_area_), as well as measurements of total nitrogen content expressed both per unit leaf mass (*N*_mass_) and per unit leaf area (*N*_area_). In addition, for each unique species observation in this dataset with a corresponding measurement for both *A*_mass_ and *N*_mass_ or for both *A*_area_ and *N*_area_, the mass-based and area-based photosynthetic nitrogen-use efficiencies were also derived using the calculations *A*_mass_/*N*_mass_ (*PNUE*_mass_) and *A*_area_/*N*_area_ (*PNUE*_area_), respectively. In cases where duplicate entries for a parameter were present across species, the mean value was taken so as to collapse the dataset to contain only a single row per species. Finally, although photosynthetic measurements were available from individuals belonging to all major land plant lineages (including the mosses, liverworts, fern allies, ferns, gymnosperms, and angiosperms), only the subset of angiosperms in the dataset for which a publicly available *rbcL* sequence could be obtained were taken forward. This is because various diffusional and biochemical factors other than rubisco are known to cause reduced photosynthetic capacities in non-angiosperm plants^85^ that would bias the results of the current study. For the same reasons, only the subset of C_3_ angiosperms in this dataset were taken forward in the present analysis to avoid picking up photosynthetic effects which result from CO_2_ concentrating mechanisms that act upstream of rubisco. In total, this left a photosynthetic dataset of 366 C_3_ angiosperms from which to base the analyses herein. Combined, this resulting dataset included 272 unique species measurements for *N*_mass_, 137 unique species measurements for *A*_mass_ and 118 unique species measurements for *PNUE*_mass_, as well as 270 unique species measurements for *N*_area_, 151 unique species measurements for *A*_area_, and 120 unique species measurements for *PNUE*_area_, respectively.

To compute the relative extents of RbcL molecular evolution which has occurred in each angiosperm in the photosynthetic dataset, the exact same method was followed as described above. First, the full RbcL phylogenetic gene tree in Supplemental File 8 that was previously inferred from the complete set of publicly available RbcL sequences in NCBI was pruned so as to contain only terminal and internal branches corresponding to angiosperms in the photosynthetic dataset. Here, in situations where duplicate sequences in the alignment resulted in multiple terminal nodes for a given species, only a single node was retained based on the sequence which is first in the alphabetical order of the gene accession numbers. As above, this reduced RbcL tree was then manually rooted in Dendroscope^140^, and the relative extent of RbcL protein evolution in each angiosperm was computed as the summed branch length (sequence substitutions per aligned sequence site) leading from the last common ancestor at the root of this clade to each respective terminal node. Finally, linear regression models were employed to assess the pairwise relationships between the variation in rubisco molecular evolution and each respective photosynthetic parameter. The resulting full integrated dataset containing photosynthetic measurements and comparable extents of RbcL molecular evolution for all 366 C_3_ angiosperms can be found in Supplemental File 9. The RbcL phylogenetic gene tree which has been pruned from that in Supplemental File 8 to contain the subset of C_3_ angiosperms in the photosynthetic dataset used for the basis of this analysis can be found in Supplemental File 10.

## Data Availability

All data used in this study is provided in the supplemental material.

## Author Contributions

JWB and SK conceived the study and wrote and edited the manuscript. JWB conducted the analysis. DME provided advice on design and implementation of the study.

