## Supplemental File 1 for "Rubisco is evolving for improved catalytic efficiency and CO_2_ assimilation in plants"

### *Form I rubisco ancillary chaperones exhibit a variable rate of molecular evolution*

The molecular evolution of interacting partners of the holoenzyme were evaluated following the same method that was used to compute the percentile rank rate of evolution in *rbcL*/RbcL and *rbcS*/RbcS. The list of interacting partners included components involved in holoenzyme metabolic regulation (RUBISCO ACTIVASE (Rca)) as well as in holoenzyme folding and assembly (CHAPERONIN 10 (Cpn10), CHAPERONIN 20 (Cpn20), CHAPERONIN-60 (Cpn60), RBCX (RbcX), RUBISCO ACCUMULATION FACTOR 1 (Raf1), RUBISCO ASSEMBLY FACTOR 2 (Raf2)). This analysis revealed that the percentile rate of molecular evolution in rubisco chaperones was highly variable across all taxonomic groups, with no consistent pattern emerging in the rate of evolution in these chaperone genes compared to that experienced by the cohort of all other genes encoded in each species genome (Figure S3A and 3B and table S11). Thus, similar to the case in the rubisco small subunit, rubisco chaperones do not experience the ubiquitous slow pace of molecular evolution which is experienced by the large subunit of the holoenzyme.

### *Correlations in rubisco molecular and kinetic evolution are robust to biases in species sampling and model of sequence evolution*

In order to evaluate the robustness of the integrated molecular and kinetic analysis we conducted a systematic and comprehensive examination of possible methodological biases associated with our investigation to determine whether these may have impacted the results which are presented. Specifically, two such potential methodological issues were identified and include biases arising from incomplete species sampling in the rubisco kinetic dataset as well as biases arising from uncertainties or errors in the underlying phylogenetic gene tree that is used as the basis of the current analysis. As such, to address each of these concerns in turn, a supporting set of additional analyses were designed, and the results generated from these analyses were compared to those generated from the original analysis to assess the reliability of the presented conclusions to potential methodological artefacts.

To account for potential methodological biases in our investigation that are associated with incomplete species sampling (including overrepresentation of certain clades on the angiosperm phylogenetic tree of life), we repeated the analysis of rubisco molecular and kinetic evolution but including only the minimum subset of species that captured the majority of phylogenetic diversity across all species in the full analysis (see Methods, Figure S4). As such, in this analysis, specifically any potential methodological artefacts caused by biases in sampling from specific clades within the tree have been addressed.

To account for potential methodological biases in our investigation that are associated with sources of error in the underlying phylogenetic tree, we repeated the analysis of rubisco molecular and kinetic evolution but using alternate RbcL phylogenetic gene trees independently inferred from the same sequence alignment but using alternate models of sequence evolution (see Methods). As such, in this analysis, any potential methodological artefacts caused by assuming a particular model of sequence evolution are removed.

Here, as analogous conclusions were generated from the results of each of these analyses (Table S11 and S12) to those presented in the main text (Figure 4B). Thus, the results presented in the main text are not an artefact caused by either systematic biases in species sampling or caused by errors in phylogenetic reconstruction.

### *Supplemental Figures*

#### Figure S1

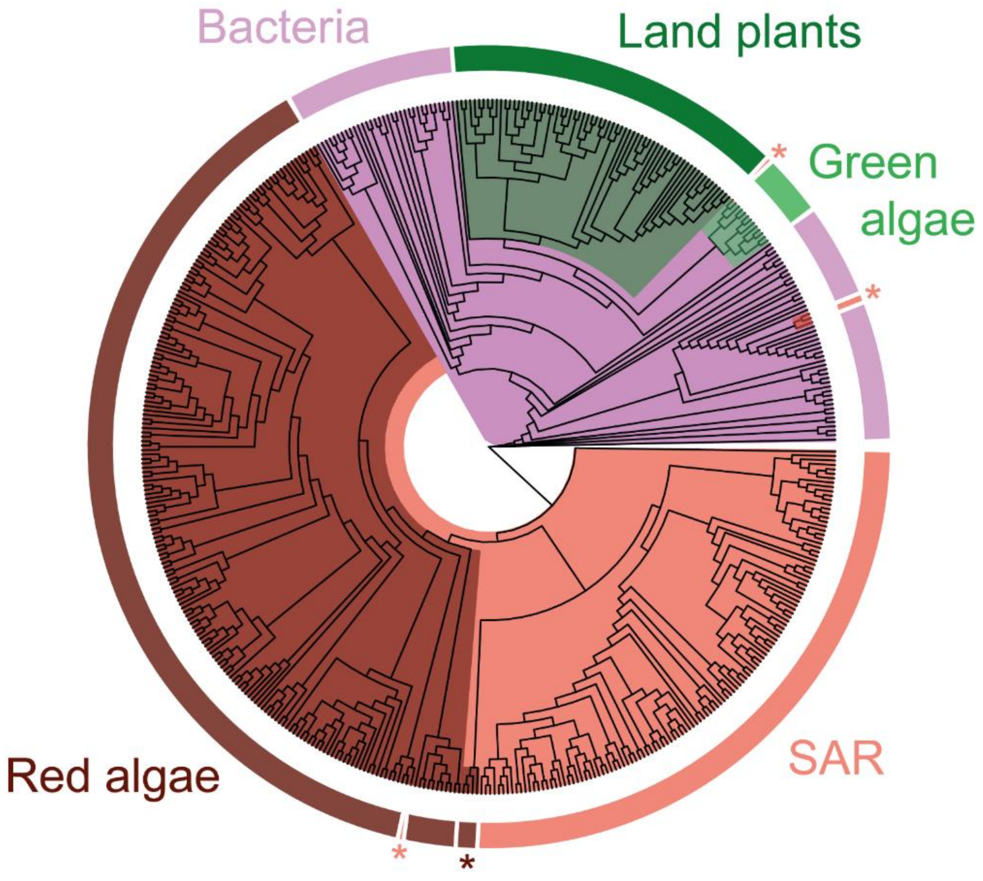


**Figure S1**. A phylogenetic tree of species in this study. Evolutionary history has been inferred from a multiple sequence alignment of the *rbcL* coding sequence in each species. The phylogeny is displayed as a cladogram for ease of visualisation, and species membership to different taxonomic groups (labelled) are highlighted by colour. Dark brown: red algae (*Rhodophyta*; *n* = 201). Light brown: SAR supergroup (*Stramenopiles*, *Alveolates*, and *Rhizaria*; *n* = 129). Lilac: bacteria (*Bacteria*; *n* = 78). Dark green: land plants (*Streptophyta*; *n* = 68). Light green: green algae (*Chlorophyta*; *n* = 12). Species at terminal nodes which cluster outside their designated taxonomic group are marked by an asterisk.

**Figure S2**

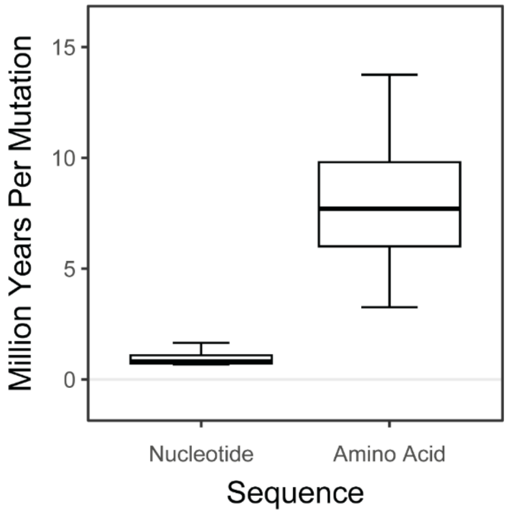


**Figure S2.** The rate of nucleotide and amino acid mutation in the rubisco large subunit experienced for $C_3$ angiosperms in the kinetic dataset, as expressed as the number of million years per individual sequence change. Molecular mutation rates were calculated since divergence from the last common ancestor at the base of the angiosperm clade 160 million years ago and considered an average *rbcL* of length 1428 nucleotide residues and 476 amino acid residues.

**Figure S3**

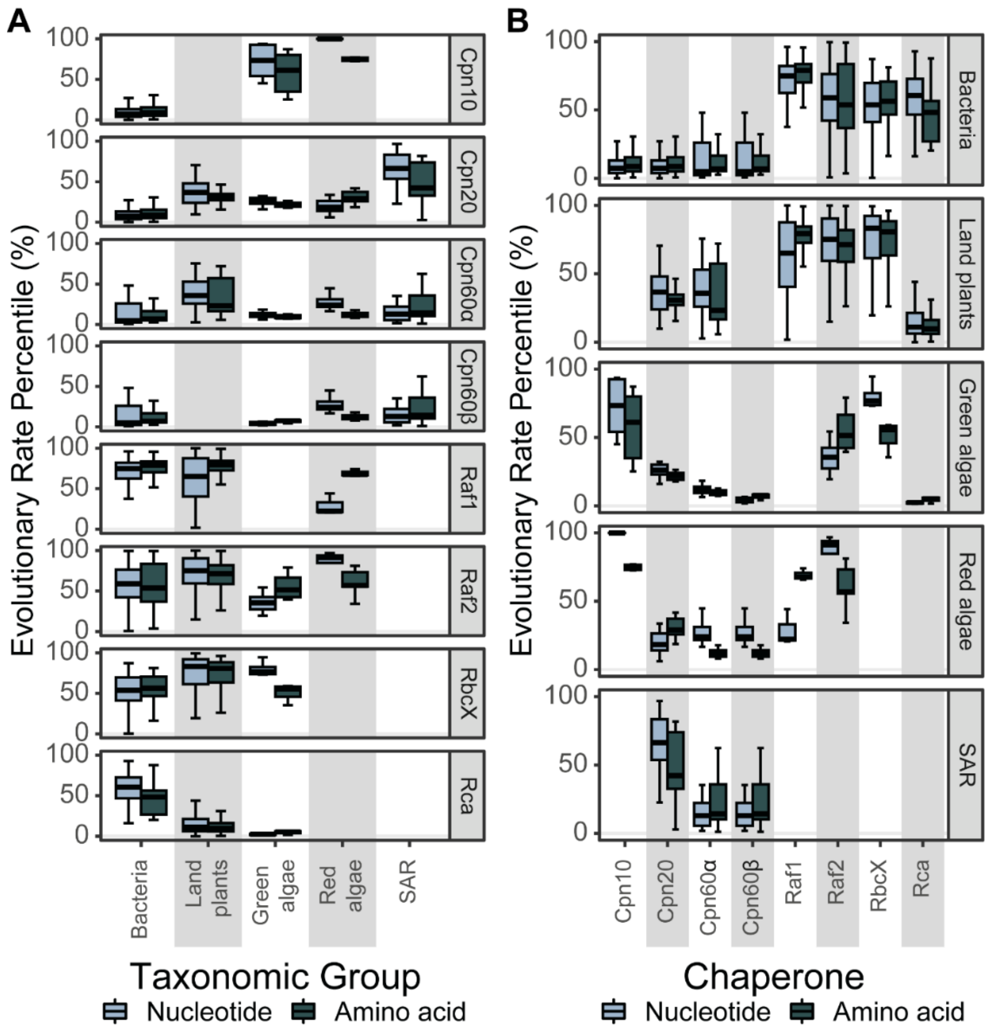


**Figure S3**. The extent of molecular evolution in each of rubisco's interacting chaperone partners. A) Boxplot of the extent of nucleotide and amino acid evolution (substitutions per sequence site) of each chaperone organised by taxonomic group. B) As in (A), but organised by gene/protein in alphabetical order. Chaperonin-60α: Cpn60α. Chaperonin -60: Cpn60β. Chaperonin -10: Cpn10. Chaperonin -20: Cpn20.RbcX: RbcX. Rubisco accumulation factor 1: Raf1. Rubisco accumulation factor 2: Raf2. Rubisco activase: Rca. Although Bundle sheath defective 2 (Bsd2) is a known regulator of rubisco assembly, this chaperone was omitted from the analysis owing to insufficient sequence data. Data for this figure can be found in Table S11.

**Figure S4**

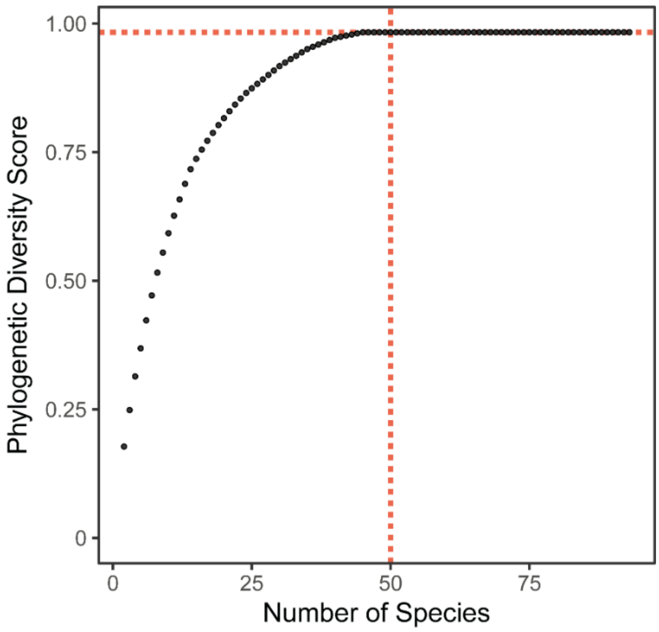


**Figure S4.** The sample size ($n$ = 50) at which phylogenetic diversity saturates is indicated by a red dashed line. The optimal species set at this threshold are shown in table S10.

### *Supplemental Tables*

#### *Table S1*

**Table S1.** Summary of the sequence dataset used for analysis of rubisco molecular evolution in this study.

| **Taxonomic Group** | ***n*** | | | |
|---|---|---|---|---|
| | **Species** | **Species Representative Gene Models** | ***rbcL*/RbcL Sequences** | ***rbcS*/RbcS Sequences** |
| Bacteria | 78 | 32 | 78 | 86 |
| Land plants | 68 | 27 | 68 | 172 |
| Green algae | 12 | 4 | 12 | 24 |
| Red algae | 201 | 6 | 201 | 595 |
| SAR | 129 | 8 | 129 | 283 |
| **Total** | **488** | **77** | **488** | **1140** |

### *Table S2*

**Table S2.** Summary of the percentile rate of nucleotide and protein evolution in each rubisco subunit in context of all other genes in each taxonomic group. The median, first quartile (Q1), third quartile (Q3), the interquartile range (IQR), the mean and the standard error (S.E.) are provided.

| **Taxonomic Group** | **Rubisco Subunit** | **Sequence** | **Percentile Rate** | | | | | |
|---|---|---|---|---|---|---|---|---|
| | | | **Median** | **Q1** | **Q3** | **IQR** | **Mean** | **S.E.** |
| Bacteria | **RbcL** | **Nucleotide** | 2.597 | 0.847 | 5.693 | 4.846 | 4.729 | 0.317 |
| Land plants | | | 0.057 | 0.016 | 0.162 | 0.146 | 0.357 | 0.100 |
| Green algae | | | 0.042 | 0.019 | 0.088 | 0.070 | 0.053 | 0.016 |
| Red algae | | | 0.043 | 0.030 | 0.192 | 0.162 | 0.132 | 0.040 |
| SAR | | | 0.575 | 0.269 | 2.005 | 1.736 | 1.653 | 0.527 |
| Bacteria | | **Protein** | 2.652 | 0.952 | 5.129 | 4.177 | 3.993 | 0.246 |
| Land plants | | | 2.180 | 1.573 | 3.314 | 1.741 | 3.092 | 0.201 |
| Green algae | | | 0.983 | 0.478 | 1.703 | 1.224 | 1.178 | 0.344 |
| Red algae | | | 0.933 | 0.874 | 1.141 | 0.267 | 1.168 | 0.186 |
| SAR | | | 2.614 | 2.048 | 3.840 | 1.792 | 3.169 | 0.467 |
| Bacteria | **RbcS** | **Nucleotide** | 25.205 | 14.899 | 39.836 | 24.937 | 28.373 | 0.837 |
| Land plants | | | 64.057 | 52.487 | 76.510 | 24.023 | 61.857 | 1.089 |
| Green algae | | | 3.961 | 1.905 | 5.643 | 3.738 | 5.198 | 2.078 |
| Red algae | | | 1.256 | 1.010 | 1.920 | 0.910 | 1.381 | 0.201 |
| SAR | | | 5.185 | 1.106 | 10.724 | 9.618 | 7.636 | 1.532 |
| Bacteria | | **Protein** | 41.927 | 24.678 | 52.779 | 28.101 | 39.030 | 0.842 |
| Land plants | | | 52.024 | 36.131 | 67.519 | 31.387 | 52.229 | 1.002 |
| Green algae | | | 14.338 | 10.363 | 17.657 | 7.293 | 14.623 | 2.020 |
| Red algae | | | 8.630 | 7.019 | 16.601 | 9.581 | 11.352 | 1.697 |
| SAR | | | 12.445 | 4.047 | 17.256 | 13.210 | 12.410 | 1.518 |

***Table S3***

**Table S3.** As in Table S2 but in context of all other enzyme-encoding genes in each taxonomic group.

| Taxonomic Group | Rubisco Subunit | Sequence | Percentile Rate | | | | | |
|---|---|---|---|---|---|---|---|---|
| | | | **Median** | **Q1** | **Q3** | **IQR** | **Mean** | **S.E.** |
| Bacteria | **RbcL** | **Nucleotide** | 3.090 | 0.993 | 7.016 | 6.023 | 5.573 | 0.365 |
| Land plants | | | 0.070 | 0.031 | 0.174 | 0.143 | 0.384 | 0.108 |
| Green algae | | | 0.057 | 0.053 | 0.061 | 0.008 | 0.057 | 0.002 |
| Red algae | | | 0.093 | 0.077 | 0.161 | 0.085 | 0.152 | 0.037 |
| SAR | | | 0.498 | 0.241 | 1.316 | 1.075 | 1.275 | 0.452 |
| Bacteria | | **Protein** | 2.978 | 0.961 | 5.706 | 4.745 | 4.334 | 0.274 |
| Land plants | | | 2.688 | 1.868 | 4.146 | 2.278 | 3.802 | 0.234 |
| Green algae | | | 0.229 | 0.192 | 0.286 | 0.094 | 0.334 | 0.115 |
| Red algae | | | 0.512 | 0.395 | 0.859 | 0.464 | 0.886 | 0.251 |
| SAR | | | 2.286 | 1.667 | 3.104 | 1.437 | 2.728 | 0.441 |
| Bacteria | **RbcS** | **Nucleotide** | 30.412 | 18.002 | 46.773 | 28.771 | 32.932 | 0.944 |
| Land plants | | | 71.256 | 60.765 | 83.175 | 22.410 | 68.478 | 1.055 |
| Green algae | | | 2.874 | 1.022 | 5.656 | 4.634 | 4.282 | 1.799 |
| Red algae | | | 1.044 | 0.525 | 2.056 | 1.531 | 1.382 | 0.299 |
| SAR | | | 6.122 | 0.819 | 11.510 | 10.692 | 8.097 | 1.639 |
| Bacteria | | **Protein** | 47.314 | 29.298 | 56.569 | 27.271 | 43.035 | 0.887 |
| Land plants | | | 61.809 | 45.382 | 76.873 | 31.491 | 60.920 | 1.016 |
| Green algae | | | 17.982 | 12.037 | 23.533 | 11.496 | 17.801 | 2.583 |
| Red algae | | | 11.933 | 8.110 | 21.443 | 13.334 | 15.126 | 2.461 |
| SAR | | | 11.738 | 3.844 | 22.623 | 18.780 | 14.133 | 1.911 |

### *Table S4*

**Table S4.** Summary of the rate of nucleotide and protein evolution in each Calvin-Benson-Bassham cycle enzyme as a percentage ratio (%) of that measured in the rubisco large subunit (*rbcL*/RbcL). RBCS: rubisco small subunit. PGK: phosphoglycerate kinase. GAPDH-A: glyceraldehyde-3-phosphate dehydrogenase A subunit. GAPDH-B: glyceraldehyde-3-phosphate dehydrogenase B subunit. TPI: triose phosphate isomerase. FBA: fructose-bisphosphate aldolase. FBP: fructose-1,6-bisphosphatase. TKL: transketolase. SBP: sedoheptulose-bisphosphatase. RPI: ribose 5-phosphate isomerase. RPE: ribulose-p-3-epimerase. PRK: phosphoribulokinase.

| **Enzyme** | **Sequence** | **% of *rbcL*/RbcL Molecular Evolution** | | | | | |
|---|---|---|---|---|---|---|---|
| | | **Median** | **Q1** | **Q3** | **IQR** | **Mean** | **S.E.** |
| RBCS | **Nucleotide** | 336.230 | 303.768 | 465.258 | 161.490 | 14215.472 | 6977.094 |
| PGK | | 194.398 | 154.133 | 260.690 | 106.557 | 10222.926 | 8653.225 |
| GAPDH-A | | 211.457 | 181.128 | 268.329 | 87.201 | 12074.030 | 9018.351 |
| GAPDH-B | | 146.379 | 112.891 | 298.844 | 185.954 | 3293.398 | 1821.487 |
| TPI | | 216.343 | 173.907 | 284.861 | 110.954 | 5679.185 | 3315.866 |
| FBA | | 221.158 | 168.605 | 302.040 | 133.435 | 10306.953 | 8831.565 |
| FBP | | 292.943 | 235.488 | 399.049 | 163.561 | 10201.657 | 5517.012 |
| TKL | | 239.675 | 202.167 | 303.532 | 101.365 | 9195.482 | 5239.744 |
| SBP | | 232.694 | 204.043 | 305.004 | 100.962 | 4667.535 | 3302.289 |
| RPI | | 426.887 | 268.135 | 694.738 | 426.602 | 15006.493 | 11123.542 |
| RPE | | 204.113 | 181.546 | 283.966 | 102.419 | 5462.525 | 4008.166 |
| PRK | | 223.418 | 159.157 | 288.776 | 129.619 | 3818.743 | 2612.882 |
| RBCS | **Protein** | 677.203 | 513.747 | 880.073 | 366.326 | 13630.350 | 6097.334 |
| PGK | | 188.897 | 128.489 | 271.151 | 142.662 | 1478.183 | 1094.706 |
| GAPDH-A | | 168.140 | 103.943 | 231.241 | 127.298 | 7539.044 | 5584.847 |
| GAPDH-B | | 102.634 | 69.460 | 180.491 | 111.031 | 7341.694 | 5148.096 |
| TPI | | 227.879 | 190.426 | 290.050 | 99.624 | 3948.902 | 2281.597 |
| FBA | | 145.765 | 107.797 | 180.354 | 72.556 | 5257.912 | 5090.570 |

| | | | | | | |
|---|---|---|---|---|---|---|
| FBP | 267.831 | 189.307 | 397.150 | 207.842 | 25149.320 | 19153.272 |
| TKL | 284.397 | 225.285 | 339.574 | 114.289 | 5429.319 | 3058.388 |
| SBP | 141.192 | 96.510 | 217.906 | 121.396 | 2784.952 | 2048.169 |
| RPI | 186.521 | 114.161 | 284.244 | 170.083 | 12075.717 | 11840.183 |
| RPE | 233.601 | 167.671 | 306.208 | 138.536 | 2686.190 | 1714.928 |
| PRK | 167.644 | 132.689 | 248.041 | 115.352 | 1892.755 | 1245.509 |

### *Table S5*

**Table S5.** One-Sample Wilcoxon Signed Rank Test to assess significant differences in the rate of nucleotide and protein evolution between the rubisco large subunit (*rbcL*/RbcL) and each Calvin-Benson-Bassham cycle enzyme. A non-parametric test was used as data failed to conform to normality (Shapiro-Wilk test; $p < 0.05$). Statistics are rounded to three decimal places and corrected significance values are represented as α levels, where; $\alpha = 0.001$ if $P < 0.001$, $\alpha = 0.01$ if $0.001 < P < 0.01$, $\alpha = 0.05$ if $0.01 < P < 0.05$, and α = ns if $P > 0.05$. Calvin-Bensen-Bassham cycle enzymes/subunits are abbreviated following the convention in table S4.

| Enzyme | Nucleotide | | Protein | |
|---|---|---|---|---|
| | Statistic | *α* | Statistic | *α* |
| RBCS | 2476425 | 0.001 | 2476417 | 0.001 |
| PGK | 51040 | 0.001 | 48937 | 0.001 |
| GAPDH-A | 55278 | 0.001 | 49267 | 0.001 |
| GAPDH-B | 2550 | 0.001 | 1689 | 0.01 |
| TPI | 59685 | 0.001 | 59570 | 0.001 |
| FBA | 8984 | 0.001 | 7727 | 0.001 |
| FBP | 60378 | 0.001 | 60261 | 0.001 |
| TKL | 60378 | 0.001 | 60378 | 0.001 |
| SBP | 59684 | 0.001 | 50188 | 0.001 |
| RPI | 37128 | 0.001 | 33896 | 0.001 |
| RPE | 45450 | 0.001 | 45030 | 0.001 |
| PRK | 50721 | 0.001 | 49118 | 0.001 |

#### *Table S6*

**Table S6.** Mean values of the ratio of rubisco large to small subunit percentile rank rate of evolution (*rbcL* to *rbcS* and RbcL to RbcS, respectively) and associated variation (±1 S.E.) in each taxonomic group.

| % Ratio | Taxonomic Group | | | | |
|---|---|---|---|---|---|
| | **Bacteria** | **Land Plants** | **Green Algae** | **Red Algae** | **SAR** |
| ***rbcL* : *rbcS*** | 30.7 ± 4.3 | 0.6 ± 0.2 | 3.1 ± 1.6 | 10.7 ± 2.8 | 70.9 ± 46.8 |
| **RbcL : RbcS** | 12.2 ± 0.7 | 8.3 ± 1.2 | 8.3 ± 2.1 | 14.7 ± 3.0 | 40.2 ± 7.2 |

***Table S7***

**Table S7.** List of all species in each taxonomic group for which either a nuclear (land plants, green algae, red algae, SAR) or bacterial (bacteria) genome could be acquired.

| Taxonomic Group | Species |
|---|---|
| **Bacteria** | Acaryochloris marina |
| | Acidithiobacillus ferrooxidans |
| | Allochromatium vinosum |
| | Anabaenopsis circularis |
| | Arthrospira platensis |
| | Aurantimonas manganoxydans |
| | Crocosphaera subtropica |
| | Gloeobacter kilaueensis |
| | Gloeobacter violaceus |
| | Gloeomargarita lithophora |
| | Halomicronema hongdechloris |
| | Hydrogenophaga pseudoflava |
| | Methylacidimicrobium cyclopophantes |
| | Methylacidimicrobium tartarophylax |
| | Methylacidiphilum fumariolicum |
| | Methylacidiphilum infernorum |
| | Microcystis aeruginosa |
| | Microcystis viridis |
| | Nocardia nova |
| | Nocardia seriolae |
| | Novimethylophilus kurashikiensis |
| | Phaeobacter gallaeciensis |
| | Phormidesmis priestleyi |
| | Planktothrix agardhii |

| | |
|---|---|
| | Prochlorococcus marinus |
| | Prochlorothrix hollandica |
| | Raphidiopsis brookii |
| | Synechococcus elongatus |
| | Thermosynechococcus elongatus |
| | Thermosynechococcus vulcanus |
| | Thioflexothrix psekupsii |
| | Trichormus variabilis |
| **Land Plants** | Aegilops tauschii |
| | Amaranthus hypochondriacus |
| | Arabidopsis thaliana |
| | Brassica napus |
| | Brassica oleracea |
| | Brassica rapa |
| | Camellia sinensis |
| | Capsicum annuum |
| | Cucumis sativus |
| | Dendrobium catenatum |
| | Glycine soja |
| | Gossypium hirsutum |
| | Hevea brasiliensis |
| | Hordeum vulgare |
| | Lactuca sativa |
| | Mucuna pruriens |
| | Nicotiana attenuata |
| | Oryza sativa |
| | Panicum virgatum |
| | Phaseolus vulgaris |

| | |
|---|---|
| | Salvia splendens |
| | Sorghum bicolor |
| | Spinacia oleracea |
| | Triticum aestivum |
| | Triticum turgidum |
| | Triticum urartu |
| | Zea mays |
| **Green Algae** | Botryococcus braunii |
| | Chromochloris zofingiensis |
| | Dunaliella salina |
| | Volvox carteri |
| **Red Algae** | Chondrus crispus |
| | Cyanidiococcus yangmingshanensis |
| | Galdieria sulphuraria |
| | Gracilariopsis chorda |
| | Porphyra umbilicalis |
| | Porphyridium purpureum |
| **SAR** | Aureococcus anophagefferens |
| | Ectocarpus siliculosus |
| | Fistulifera solaris |
| | Microchloropsis salina |
| | Nannochloropsis gaditana |
| | Phaeodactylum tricornutum |
| | Thalassiosira oceanica |
| | Thalassiosira pseudonana |

### *Table S8*

**Table S8.** The gene loci encoding the photosynthetic isoforms of Calvin-Benson-Bassham cycle enzymes in *Arabidopsis thaliana*.

| Enzyme | Unique ID | Gene name | Arabidopsis TAIR ID |
|---|---|---|---|
| PGK | PGK1 | PHOSPHOGLYCERATE KINASE 1 | AT3G12780 |
| | PGK2 | PHOSPHOGLYCERATE KINASE 2 | AT1G56190 |
| GAPDH-A | GAPA-1 | GLYCERALDEHYDE 3-PHOSPHATE DEHYDROGENASE A SUBUNIT 1 | AT3G26650 |
| | GAPA-2 | GLYCERALDEHYDE 3-PHOSPHATE DEHYDROGENASE A SUBUNIT 2 | AT1G12900 |
| GAPDH-B | GAPB | GLYCERALDEHYDE-3-PHOSPHATE DEHYDROGENASE B SUBUNIT | AT1G42970 |
| TPI | TPI | TRIOSEPHOSPHATE ISOMERASE | AT2G21170 |
| FBA | FBA1 | FRUCTOSE-BISPHOSPHATE ALDOLASE 1 | AT2G21330 |
| | FBA2 | FRUCTOSE-BISPHOSPHATE ALDOLASE 2 | AT4G38970 |
| FBP | FBP1 | FRUCTOSE 1,6-BISPHOSPHATE PHOSPHATASE | AT3G54050 |
| TKL | TKL1 | TRANSKETOLASE 1 | AT3G60750 |
| SBP | SBP | SEDOHEPTULOSE-BISPHOSPHATAS | AT3G55800 |
| RPI | RPI | RIBOSE 5-PHOSPHATE ISOMERASE | AT3G04790 |
| RPE | RPE | D-RIBULOSE-5-PHOSPHATE-3-EPIMERASE | AT5G61410 |
| PRK | PRK | PHOSPHORIBULOKINASE | AT1G32060 |

***Table S9***

**Table S9.** The gene loci encoding the chaperones involved in Form I rubisco assembly and metabolic regulation in *Arabidopsis thaliana*.

| Chaperone | Unique ID | Gene name | Arabidopsis TAIR ID |
|---|---|---|---|
| Bsd2 | Bsd2 | BUNDLE SHEATH DEFECTIVE 2 | AT3G47650 |
| Cpn60α | Cpn60α_1 | CHAPERONIN-60ALPHA1 | AT2G28000 |
| | Cpn60α_2 | CHAPERONIN-60ALPHA2 | AT5G18820 |
| Cpn60β | Cpn60β_1 | CHAPERONIN-60BETA1 | AT1G55490 |
| | Cpn60β_2 | CHAPERONIN-60BETA2 | AT3G13470 |
| | Cpn60β_3 | CHAPERONIN-60BETA3 | AT5G56500 |
| | Cpn60β_4 | CHAPERONIN-60BETA4 | AT1G26230 |
| Cpn10 | Cpn10_1 | CHLOROPLAST CHAPERONIN 10 | AT2G44650 |
| | Cpn10_2 | GROES | AT3G60210 |
| Cpn20 | Cpn20 | CHAPERONIN 20 | AT5G20720 |
| RbcX | RbcX_1 | RBCX1 | AT4G04330 |
| | RbcX_2 | RBCX2 | AT5G19855 |
| Raf1 | Raf1_1 | RUBISCO ACCUMULATION FACTOR-LIKE PROTEIN | AT5G28500 |
| | Raf1_2 | RUBISCO ACCUMULATION FACTOR 1 | AT3G04550 |
| Raf2 | Raf2 | RUBISCO ASSEMBLY FACTOR 2 | AT5G51110 |
| Rca | Rca | RUBISCO ACTIVASE | AT2G39730 |

***Table S10***

**Table S10.** The subset of 50 $C_3$ angiosperms which captured the vast majority of phylogenetic diversity encapsulated in the full rubisco kinetic dataset, as used for the basis of the analysis in table S11.

| **Division** | **Species** |
|---|---|
| **Dicot** | Agriophyllum squarrosum |
| | Amphicarpaea bracteata |
| | Artemisia myriantha |
| | Artemisia vulgaris |
| | Beta vulgaris |
| | Chenopodiastrum murale |
| | Chenopodium album |
| | Citrullus ecirrhosus |
| | Desmodium cinereum |
| | Desmodium intortum |
| | Erythrina flabelliformis |
| | Euphorbia helioscopia |
| | Euphorbia microsphaera |
| | Flaveria cronquistii |
| | Flaveria pringlei |
| | Flueggea suffruticosa |
| | Foeniculum vulgare |
| | Glycine canescens |
| | Lepidium campestre |
| | Macrotyloma uniflorum |
| | Manihot esculenta |
| | Mercurialis annua |
| | Nicotiana tabacum |

| | |
|---|---|
| | Oxybasis rubra |
| | Phaseolus coccineus |
| | Pueraria montana |
| | Sphenostylis stenocarpa |
| | Spinacia oleracea |
| | Tephrosia candida |
| | Tephrosia purpurea |
| | Tephrosia rhodesica |
| **Monoocot** | Aegilops comosa |
| | Aegilops speltoides |
| | Agrostis stolonifera |
| | Arctagrostis latifolia |
| | Brachypodium distachyon |
| | Bromus anomalus |
| | Calamagrostis stricta subsp. inexpansa |
| | Deschampsia danthonioides |
| | Elymus farctus |
| | Lolium giganteum |
| | Lolium multiflorum |
| | Musa velutina |
| | Oryza barthii x Oryza glaberrima |
| | Oryza eichingeri |
| | Oryza glaberrima |
| | Panicum milioides |
| | Poa palustris |
| | Puccinellia distans |
| | Puccinellia maritima |

### *Table S11*

**Table S11.** The pairwise correlation coefficient (% variation explained), associated significance and direction of association from the results of the linear regression analysis between the extent of RbcL protein evolution and each rubisco kinetic trait but when considering the minimum combination of species which optimize phylogenetic diversity of species in the kinetic dataset, as predicated on Figure S4. Significance values are represented as α levels, where α = 0.001 if P < 0.001, α = 0.01 if 0.001 < P < 0.01, α = 0.05 if 0.01 < P < 0.05, and α = ns if P > 0.05.

| Kinetic Trait | % Var Explained | *α* | Direction |
|---|---|---|---|
| $S_{C/O}$ | 13.5 | 0.01 | + |
| $k_{catC}$ | 7.5 | 0.05 | + |
| $K_{catC}/k_{C}$ | 6.4 | 0.05 | + |
| $K_{C}$ | ns | ns | ns |
| $K_{C}^{air}$ | ns | ns | ns |
| $K_{O}$ | ns | ns | ns |
| $K_{C}/K_{O}$ | ns | ns | ns |

### *Table S12*

**Table S12.** The pairwise correlation coefficient (% variation explained), associated significance and direction of association from the results of the linear regression analysis between the extent of RbcL protein evolution and each rubisco kinetic trait, but from the analysis of independent phylogenetic tree inferred from the same set of species sequence data but using an alternative model of sequence evolution. Significance values are represented as α levels, where α = 0.001 if P < 0.001, α = 0.01 if 0.001 < P < 0.01, α = 0.05 if 0.01 < P < 0.05, and α = ns if P > 0.05.

| Tree Model | Kinetic Trait | % Var Explained | *α* | Direction |
|---|---|---|---|---|
| **LG+I+G4** | $S_{C/O}$ | 7.2 | 0.01 | + |
| | $k_{catC}$ | 11.0 | 0.001 | + |
| | $K_{catC}/k_C$ | 9.6 | 0.01 | + |
| | $K_C$ | ns | ns | ns |
| | $K_C^{air}$ | ns | ns | ns |
| | $K_O$ | ns | ns | ns |
| | $K_C/K_O$ | ns | ns | ns |
| **JTTDCMut+I+G4** | $S_{C/O}$ | 7.5 | 0.01 | + |
| | $k_{catC}$ | 5.1 | 0.05 | + |
| | $K_{catC}/k_C$ | 12.3 | 0.001 | + |
| | $K_C$ | ns | ns | ns |
| | $K_C^{air}$ | ns | ns | ns |
| | $K_O$ | ns | ns | ns |
| | $K_C/K_O$ | ns | ns | ns |
